# Generation and characterization of CRISPR-Cas9-Mediated *XPC* Gene Knockout in Human Skin Cells

**DOI:** 10.1101/2024.01.25.577199

**Authors:** Ali Nasrallah, Hamid-Reza Rezvani, Farah Kobaisi, Ahmad Hammoud, Jérôme Rambert, Jos P.H. Smits, Eric Sulpice, Walid Rachidi

## Abstract

Xeroderma pigmentosum group C (XPC) is a versatile protein, crucial for sensing DNA damage in the global genome nucleotide excision repair (GG-NER) pathway. This pathway is vital for mammalian cells, acting as their essential approach for repairing DNA lesions stemming from interactions with environmental factors, such as exposure to ultraviolet (UV) radiation from the sun. Loss-of-function mutations in the *XPC* gene confer a photosensitive phenotype in XP-C patients with the accumulation of unrepaired UV induced DNA damage. This remarkable increase in DNA damage tends to elevate by 10,000-fold the risk of developing melanoma and non-melanoma skin cancers. To date, creating accurate and reproducible models to study human XP-C disease has been an important challenge. To tackle this, we used CRISPR-Cas9 technology in order to knockout *XPC* gene in various human skin cells (keratinocytes, fibroblasts, and melanocytes). After validation of the *XPC* knockout in these edited skin cells, we showed that they recapitulate the major phenotypes of XPC mutations: photosensitivity and the impairment of UV induced DNA damage repair. Moreover, these mutated cells demonstrated a reduced proliferative capacity compared to their respective wild-type controls. Finally, to better mimic the disease environment, we built a 3D reconstructed skin using these XPC knockout skin cells. This model exhibited an abnormal behavior, showing an extensive remodeling of its extracellular matrix compared to normal skin. Analyzing the composition of the fibroblasts secretome revealed a significant augmented shift in the inflammatory response following XPC knockout. Our innovative “disease on a dish” approach can provide valuable insights into the molecular mechanisms underlying XP-C disease, paving the way to design novel preventive and therapeutic strategies to alleviate the disease phenotype. Also, given the high risk of skin cancer onset in XP-C disease, our new approach can also serve as a link to draw novel insights towards this elusive field.

## Introduction

The skin, making up about 15% to 20% of an adult’s body weight and covering a surface area ranging from 1.5 to 2 m^2^, stands out as the largest organ in the human body. It fulfills numerous crucial roles, including vitamin D and melanin synthesis, sensory perception, hydration prevention, temperature regulation, acting as a primary barrier against external pathogens, and providing protection against mechanical stress^1^. Nevertheless, skin cells remain prone to metabolic disruptions because they are directly exposed to external factors. Among them, ultraviolet radiation (UVR) stands out for its potent genotoxic effects, leading to DNA damage and potentially tumor development. According to its wavelength, UVR from the sunlight can be divided into three types: UVA (λ=320–400 nm), UVB (λ=280–320 nm), and UVC (λ=100–280 nm). UVC, being absorbed by the ozone layer, leav UVA and UVB as the main sources of UV-induced DNA lesions^2^. DNA can directly absorb UVB irradiation with wavelengths between 280 and 320nm to yield dimers between adjacent pyrimidine residues. These lesions can manifest either as dewar isomers, cyclobutane pyrimidine dimers (CPDs), or 6-4 pyrimidine-pyrimidone photoproducts (6-4PPs) depending on the amount of energy absorbed by the DNA chain’s base pairs^3^. Among these, 6-4PPs induce a pronounced helical distortion in the DNA, making them readily detectable and subject to faster repair kinetics compared to CPDs^4^. CPDs, on the other hand, typically cause a weaker helical distortion in the DNA, rendering them more challenging to repair^4^. The ongoing presence of these dimers can ultimately result in the creation of double-strand breaks, as a result of replication forks collapsing. UVB irradiation predominantly leads to CC→TT or C→T transitions in DNA, and these mutations are classified as a signature for such type of irradiation^5^. Additionally, exposure to UVB radiation can generate reactive oxygen species (ROS), though at diminished levels compared to UVA irradiation, necessitating prompt repair intervention^6^.

DNA repair systems have developed throughout biological evolution to address various types of DNA damage with specificity. Upon detecting a lesion, repair systems initiate a cascade of events involving damage sensing, verification, error correction, and restoration of the initial genetic information. There are several DNA repair pathways, including nucleotide excision repair (NER), base excision repair (BER), mismatch-mediated repair (MMR), double-stranded break (DSB) repair, and others. Among these, the nucleotide excision repair (NER) pathway plays a pivotal role in removing photoproducts induced by UV light. NER process is intricate and involves a consortium of more than 40 proteins working sequentially to remove DNA damage. This DNA repair pathway can be subdivided into two main mechanisms: transcription-coupled repair (TCR) and global genome repair (GGR). TCR primarily operates in regions of actively transcribed genes, while GGR is responsible for repairing DNA damage throughout the entire genome, including both transcribed and non-transcribed DNA strands in active and dormant genes. TCR and GGR involve different recognition mechanisms for DNA damage. In GGR, a complex composed of XPC-Rad23B-Centrin2 and XPE-DDB1 is required for damage identification. XPC is adept at sensing 6-4PP lesions, while CPDs require the involvement of XPE-DDB1. TCR recognition, on the other hand, relies on CSA and CSB proteins. The subsequent repair procedures are identical for both GGR and TCR. This involves enlisting the XPD and XPB helicase components from the TFIIH complex to uncoil the DNA in the vicinity of the damaged site. Once XPA has confirmed the damage, the nucleases XPF and XPG remove the damaged displaced strand, resulting in a gap that is filled and sealed by the DNA polymerase and ligase machinery^7^.

Deficiencies in NER are linked to various disorders, including Xeroderma pigmentosum (XP), Cockayne syndrome (CS), Trichothiodystrophy (TTD), Cerebro-oculo-facio-skeletal syndrome (COFS), UV-sensitive syndrome (UVsS), and combined phenotypes, e.g. XP-CS, XP-TTD. Xeroderma Pigmentosum (XP) arising from GG-NER deficiency and can be defined as a hereditary autosomal recessive genetic disease characterized by abnormal pigmentation and an extremely hypersensitive phenotype to sunlight. XP encompasses a variable nomenclature based on the type of mutation affecting one of eight different XP genes (XPA to XPG and XPV). As a result, there is a high predisposition for cutaneous cancer onset on body parts exposed to sunlight^8^. Furthermore, internal cancers also can develop (lung, glioma, leukemia, prostate, uterus, or breast)^9^. Being the most commonly affected genetic variant in XP genodermatosis, XP-C disease, also known as Xeroderma Pigmentosum complementation group C (OMIM# 278,720), results mainly from nonsense mutations in the XPC protein, being crucial for initiating the GG-NER pathway by detecting and binding to DNA helical distortions opposite to the photoproducts generated by UV radiation^10^. Individuals with XP-C disease experience extreme photosensitivity and an accumulation of UV-induced DNA damage due to these genetic mutations. This condition is often referred to as a skin cancer-prone disease because individuals with XP-C have a significantly higher risk of developing melanomas and non-melanoma skin cancers (NMSCs) at a young age compared to healthy individuals. In fact, their susceptibility to these skin cancers is estimated to be 2,000 to 10,000 times greater than that of normal individuals^11^. This high fold risk underscores the critical role of the DNA damage recognition protein in protecting against the harmful effects of UV radiation. It’s important to note that, at present, there is no cure for XP-C syndrome. The primary approach to managing this condition involves preventive measures, such as the use of specialized UV protective shields, antioxidant creams, and sunscreens^12^. In addition to its role in Nucleotide Excision Repair (NER), XPC plays a pivotal role in various other cellular processes and DNA repair pathways. For instance, XPC is actively involved in the initial step of Base Excision Repair (BER), particularly in the removal of oxidative DNA damage^13^. Additionally, XPC plays a crucial role in maintaining the balance of cellular redox levels. Rezvani and colleagues illustrated that when XPC is deficient, it leads to the buildup of DNA damage, which, in turn, initiates the activation of AKT1. AKT1 is a well-known factor that triggers the activation of NADPH oxidase 1 (NOX1), an enzyme responsible for producing reactive oxygen species (ROS)^14^. Furthermore, Liu and their team have demonstrated the significance of XPC in apoptotic processes using zebrafish model^15^, and Magnaldo and his associates have highlighted XPC’s involvement in disturbing skin differentiation^16^.

Although some aspects regarding XPC’s function have been elucidated, there are still remain many mysteries to uncover, particularly in the context of diseased state. Lack of reliable human skin cell models for XP-C disease, along with a strong mirror control, necessitates their creation. This is essential for investigating the disease phenotype, such as molecular perturbations and cellular transformations. Various research groups have utilized primary human XP-C mutated keratinocytes and fibroblasts from patients^17,18^. However, the challenge lies in the fact that XP-C disease affects the entire body, and the absence of a wild-type control group with a similar genetic background makes it difficult to study several molecular aspects directly associated with XP-C disease. Additionally, the use of XP-C mutated fibroblasts from mice is also hindered by the differences in genetic makeup and skin architecture with humans^19^, presenting another obstacle. To overcome these barriers, precise genome editing would be of main interest to generate a reproducible XP-C disease model.

The emergence of CRISPR-Cas9 in 2012 revolutionized molecular genetics and has since become an indispensable tool for precise genetic modifications. The CRISPR-Cas9 system consists of two key components: a DNA nuclease known as Cas9, often referred to as the “molecular scissors”, and a single-stranded guide RNA (sgRNA). This sgRNA is specifically designed to guide Cas9 to particular DNA sequences of interest. By introducing the Cas9 protein transiently along with customized gRNAs that bind to target DNA sequences, a ribonucleoprotein complex (RNP) is formed. This complex can induce double-stranded breaks at precise locations in the DNA. In the CRISPR-Cas9 genome editing process, the cell’s DNA repair mechanism comes into play to repair the DNA break created by Cas9. This repair mechanism, known as non-homologous end joining (NHEJ), often leads to deletions or insertions in the DNA, resulting in gene disruption^20^. To perform functional genetics using CRISPR, specific sgRNAs are designed to target the gene(s) of interest and are produced alongside the Cas9 protein. When successful, these sgRNAs induce mutations in the target gene(s), generating loss-of-function alleles that can be further characterized after genome editing.

Taking advantage from this versatile tool, our research focused on knocking out the *XPC* gene in various human immortalized skin cells, including keratinocytes, fibroblasts, and melanocytes, using the RNP strategy. After achieving a high efficacy in editing the *XPC* gene, the knockout was validated through diverse analyses, including immunofluorescence, RT-qPCR, western blot, and Sanger sequencing. Characterizing the XP-C disease phenotype involved the photosensitivity of XPC knockout cells following UVB irradiation in a dose- and time-dependent manner (24, 48, and 72 hours). Additionally, the NER repair system impairment was quantified by analyzing 6-4PPs DNA damage 24 hours post-UVB irradiation using single-cell analysis. Moreover, the impact of XPC knockout on partially halting skin cells proliferation was elucidated. In order to accurately replicate the specific characteristics of this disease, we employed XPC knockout keratinocytes, fibroblasts, and melanocytes to construct a 3D reconstructed skin model. This model displayed an unusual profile, marked by significant degradation of its extracellular matrix in scaffold compared to the wild-type. To gain preliminary insights, we zoomed on analyzing a bunch of inflammatory markers present in the secretome of XPC knockout fibroblasts versus their associated wild-type cells, being the major cells involved influencing the skin microenvironment^21^ to decipher the consequence underlying such degradation reflecting a strong shift in the inflammatory profile. Beyond establishing a robust XPC knockout model using human skin cells, we demonstrated a straightforward, cost-effective, and efficient method for modifying skin cells, with potential applications for editing other genes of interest.

## Results

### Generation of a Complete XPC CRISPR-Cas9 KO in Human Keratinocyte, Fibroblast and Melanocyte Cell Lines

Several methods exist for introducing foreign genetic material into cells, with electroporation being one such approach^22^. In this study, we examined the editing efficiency of human immortalized skin keratinocytes (N/TERT-2G) by electroporating them with a ribonucleoprotein complex consisting of Cas9 protein and sgRNA targeting the *XPC* gene at the exon three site. This procedure resulted in a highly effective editing outcome (∼99%) in the edited heterogeneous cell population when compared to the wild-type cells (Figure 1A). The predominant form of frameshift mutation observed was deletion, with a distribution percentage of approximately two-nucleotide deletions (∼98%), one-nucleotide deletions (∼1%), and unedited sequences (∼1%) (Figure 1B). After observing the substantial editing effectiveness in the human immortalized keratinocyte cell line (N/TERT-2G), our interest extended to modeling XP-C disease in human immortalized fibroblast (S1F/TERT-1) and melanocyte (Mel-ST) cell lines so that we will have three different skin cell types from human origin and all being generated from the same immortalization strategy. This involved employing the same strategy and target site for the *XPC* gene, as illustrated (Figure 2).

**Figure 1.**
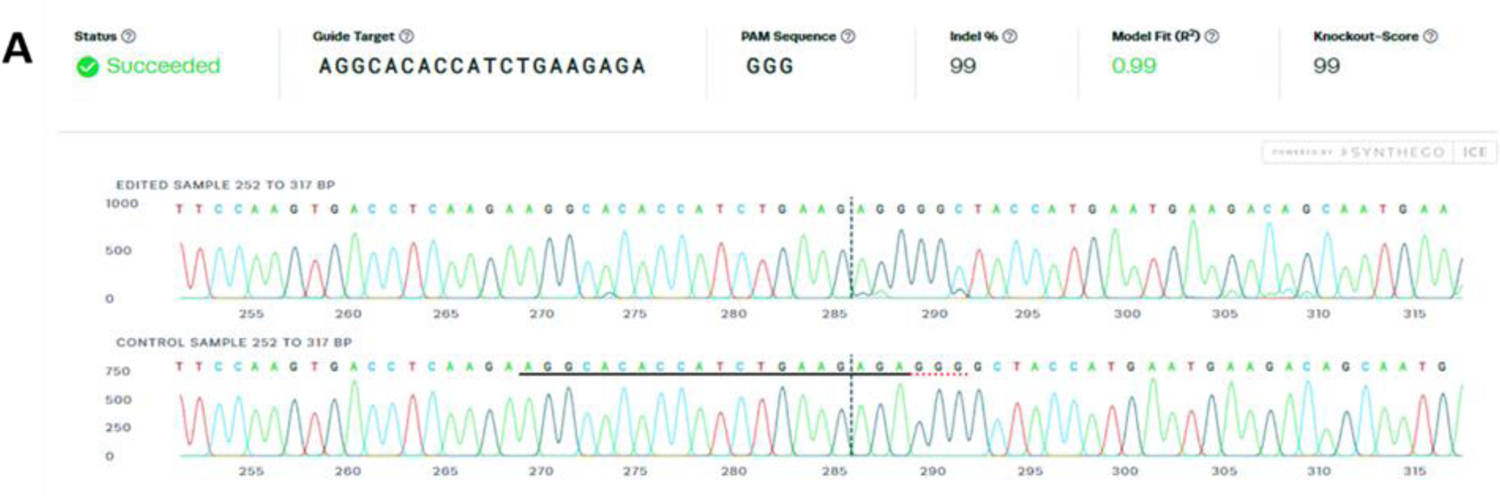
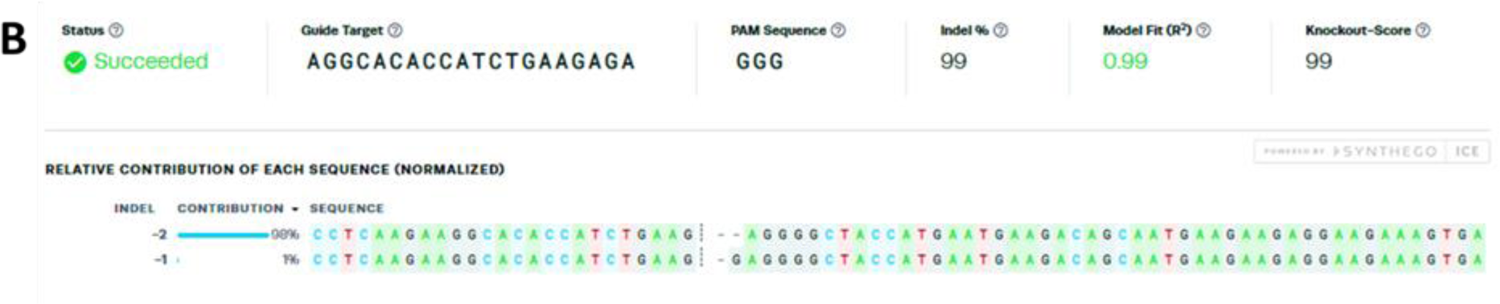
Sequencing analysis of N/TERT-2G XPC knockout (KO) heterogeneous population compared to wildtype. (A,B) The N/TERT-2G XPC knockout (KO) heterogeneous population subjected to Sanger sequencing and compared to the wild-type DNA sequence. (A) The sgRNA target region, the PAM sequence, and the knockout score are highlighted. Comparison of both sequences showed a predominant two-nucleotide (AG) indel mutation, in the exon 3 site. (B) The percentage of distribution of edits (indels) in the DNA sequence of the N/TERT-2G XPC knockout (KO) heterogeneous population.

**Figure 2.**
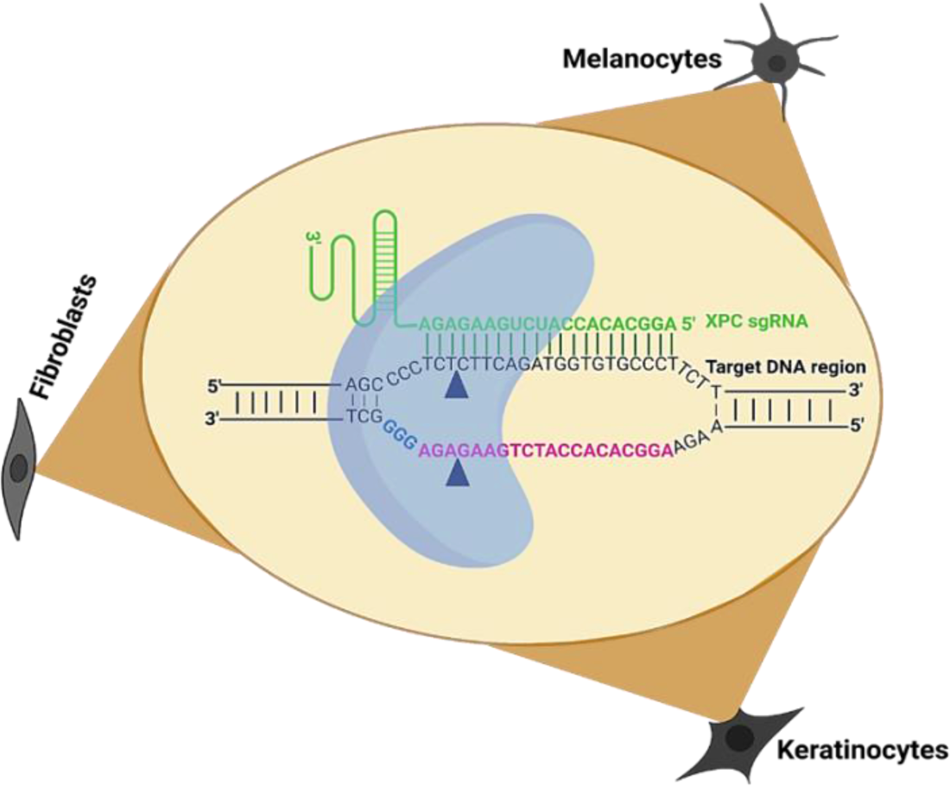
Schematic representation of the target site of *XPC* gene to be edited in human immortalized keratinocytes, fibroblasts and melanocytes.

A full homozygous knockout (KO) mutation of the *XPC* gene was successfully accomplished in human immortalized keratinocyte (N/TERT-2G), fibroblast (S1F/TERT-1), and melanocyte (Mel-ST) cell lines, employing the identical RNP strategy and the same sgRNA targeting exon three of the *XPC* gene. The edited heterogeneous populations of the three cell types underwent clonal expansion by either utilizing BD FACSMelody™ Cell Sorter or standard limiting serial dilution method to deposit a single cell in a 96-well plate. The individual single-cell clones were identified, monitored, and subsequently cultured for a period of 2 weeks for expansion. Seven single clones from each cell type were stained with XPC antibody to detect the clones with a knockout. The quantification of fluorescence corresponding to the XPC protein expression level was carried out at the single cell level. For keratinocytes, five (clones 2,4,5,6, and 7) out of seven clones were knockout compared to the wild-type control (Figure 3A). Six for fibroblasts (1,2,3,4,6, and 7) and melanocytes (1,2,3,4,5, and 7) out of seven clones were having no XPC expression level compared to their wild-type controls (Figures 3B and 4C).

**Figure 3.**
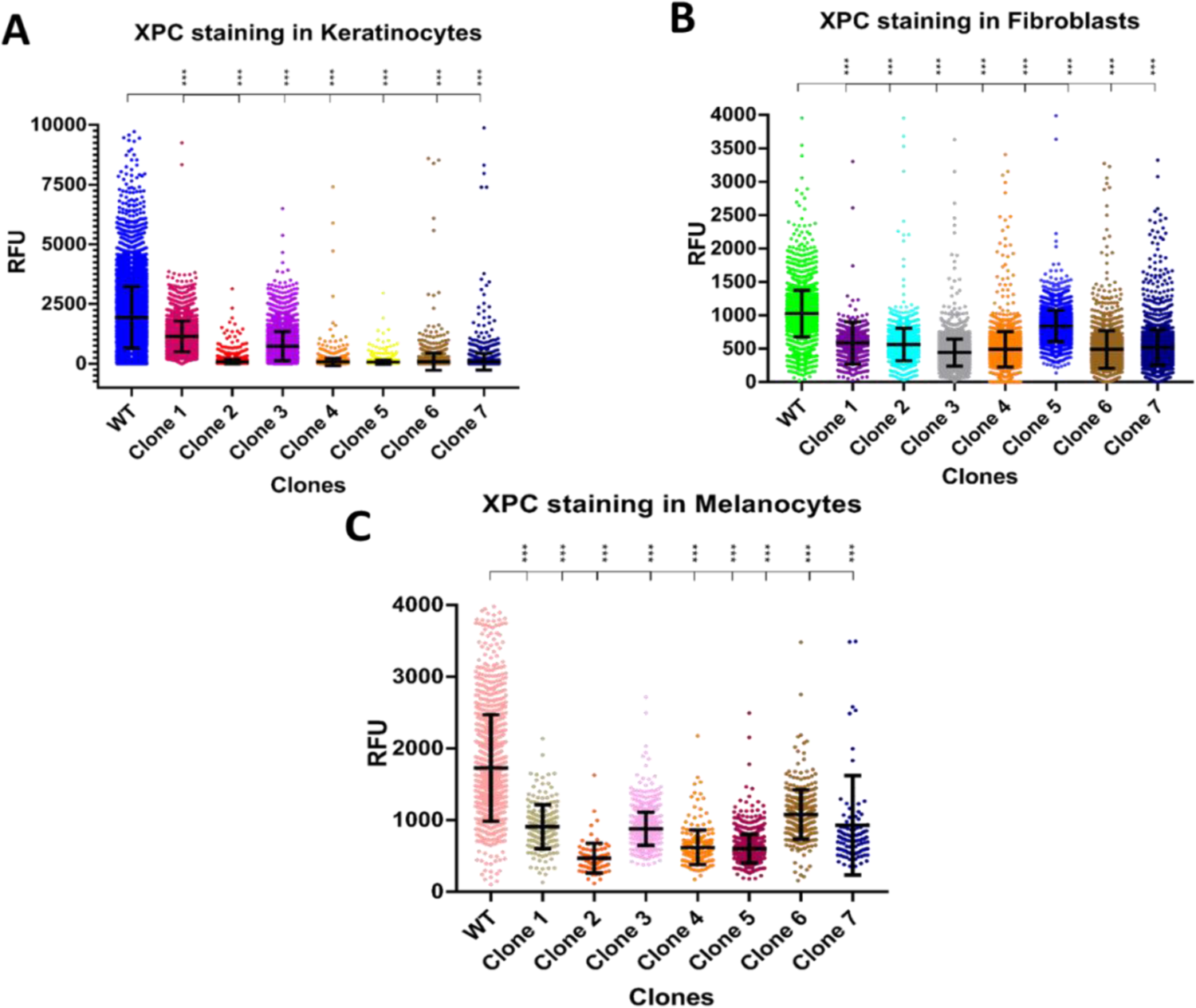
(A,B,C) Selection of the *XPC* gene homozygous knockout (KO) clones in N/TERT-2G, S1F/TERT-1 and Mel-ST cell lines. (A) Selection of the XPC gene homozygous knockout (KO) clones in N/TERT-2G cell line. Five edited N/TERT-2G clones (2,4,5,6, and 7) showed an absence of XPC’s relative fluorescence unit (RFU) compared to the wild-type (WT) control. (B) Selection of the XPC gene homozygous knockout (KO) clones in S1F/TERT-1 cell line. Six edited S1F/TERT-1 clones (1,2,3,4,6, and 7) showed an absence of XPC’s relative fluorescence unit (RFU) compared to the wild-type (WT) control. (C) Selection of the XPC gene homozygous knockout (KO) clones in Mel-ST cell line. Six edited Mel-ST clones (1,2,3,4,5, and 7) showed an absence of XPC’s relative fluorescence unit (RFU) compared to the wild-type (WT) control. *** p-value <0.001. Student T test.

**Figure 4.**
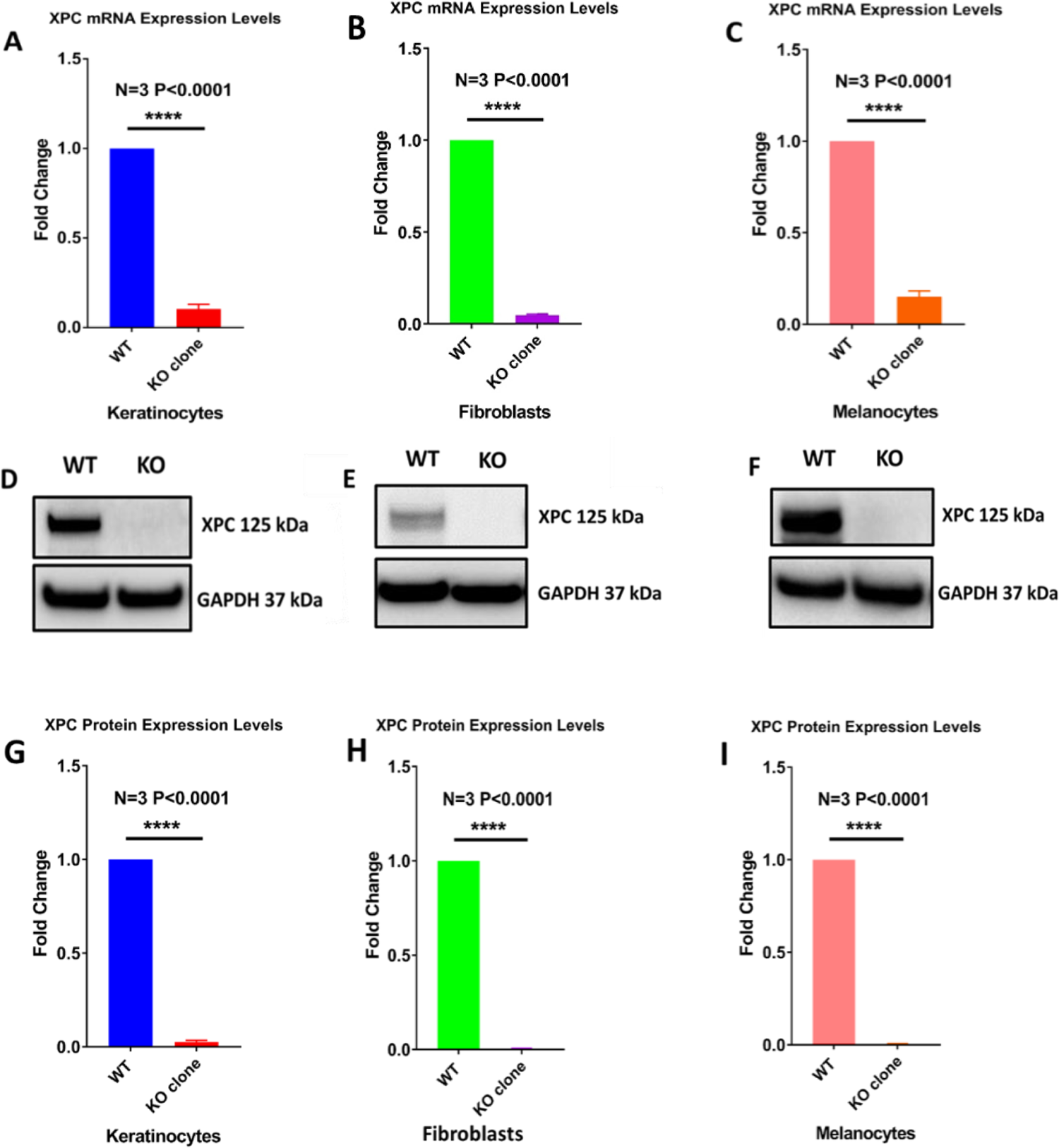
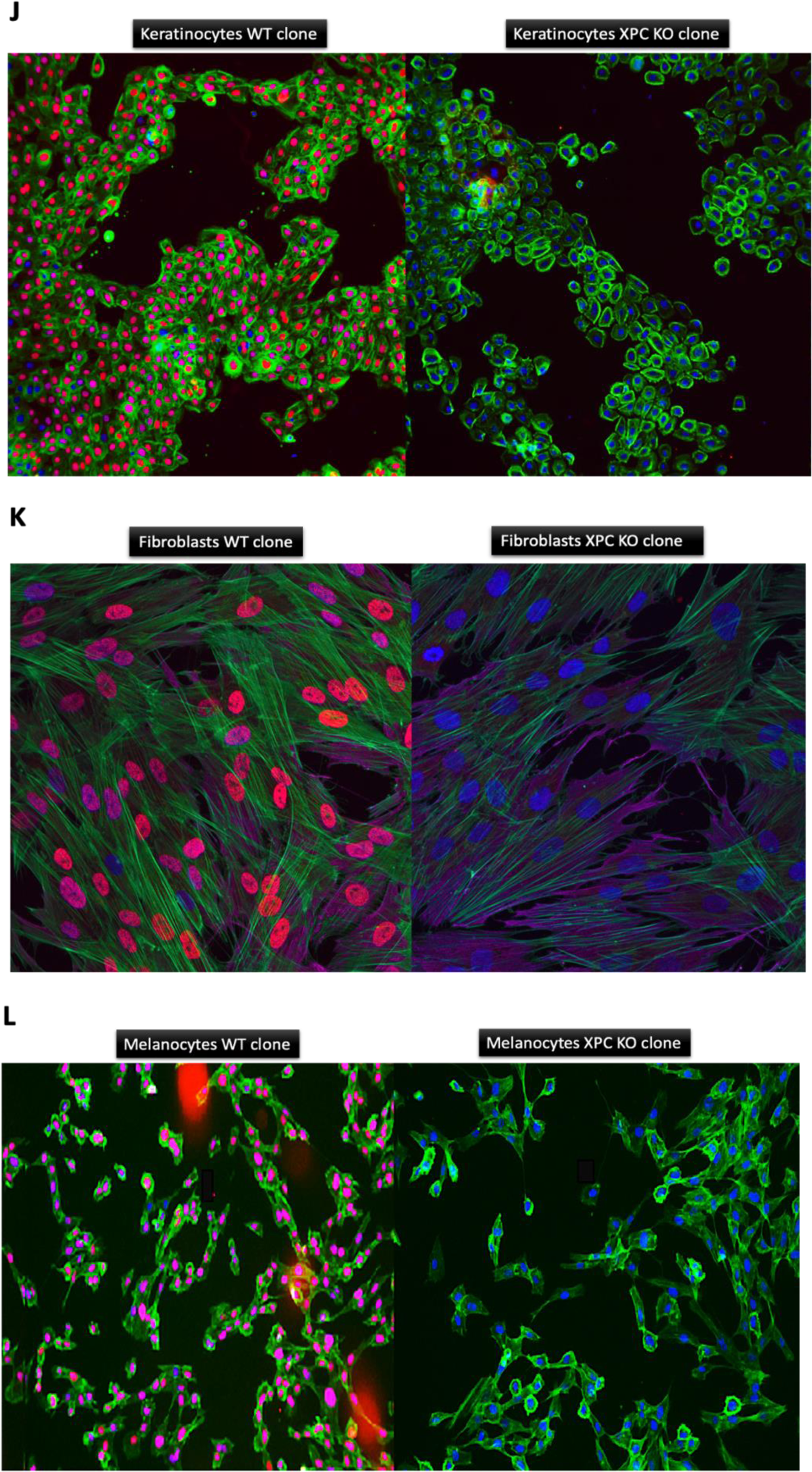
Validation of *XPC* gene knockout (KO) in N/TERT-2G, S1F/TERT-1 and Mel-ST cell lines. The expression of XPC at the mRNA and protein level in keratinocyte, fibroblast, and melanocyte cell lines was examined. RT-qPCR analysis shows an absence of XPC’s mRNA expression levels in keratinocytes (A), fibroblasts (B), and melanocytes (C) knockout clones compared to their associated wild-type cells (****p-value<0.0001) unpaired t-test. Total mRNA content was extracted and reverse transcribed into cDNAs to further quantify the expression of XPC via qPCR. Western blot analysis shows an absence of XPC’s protein expression levels in keratinocytes (D,G), fibroblasts (E,H), and melanocytes (F,I) knockout clones compared to their associated wild-type cells (****p-value<0.0001) unpaired t-test. Cellular protein extracts were separated by SDS polyacrylamide gel electrophoresis (PAGE) and transferred onto a nitrocellulose membrane. The presence or absence of XPC (band size 125 kDa), and the housekeeping protein GAPDH (band size 37 kDa) was visualized using specific antibodies (GAPDH as control). Immunofluorescence stain shows the absence of XPC protein in the nucleus of keratinocytes (J), fibroblasts (K), and melanocytes (L) knockout clones (on the right side) compared to that of their associated wild-type cells (on the left side). All cell types were stained with primary XPC antibody (cy3 in red-purple), phalloidin for the cytoplasm (green), and primary vimentin antibody for fibroblasts (cy5 in light purple). Hoechst was also used to stain the nucleus (in blue). With either 4X or 10X magnification, Image acquisition was done using Cell-insight NXT.

These results reflect the robustness of the electroporation system when combined with the ribonucleoprotein complex editing strategy to knockout the *XPC* gene. One XPC knockout (KO) clone from each cell type underwent further selection to confirm the absence of XPC mRNA and protein expression levels. RT-qPCR analyses revealed nearly absent XPC mRNA expression levels in N/TERT-2G (Figure 4A), S1F/TERT-1 (Figure 4B), and Mel-ST (Figure 4C) knockout clones when compared to their respective wild-type cells. Additionally, Immunofluorescence and western blot analyses for XPC protein (with a band size of 125 kDa) demonstrated the absence of expression in N/TERT-2G (Figures 4D, 4G, and 4J), S1F/TERT-1 (Figures 4E, 4H, and 4K), and Mel-ST (Figures 4F, 4I, and 4L) knockout clones when compared to their respective wild-type cells.

### Characterization of Wild Type and XPC KO Keratinocyte, Fibroblast and Melanocyte cell lines

#### Photosensitivity

Based on the literature, XP-C mutated cells display an elevated sensitivity to UVB radiation stemming from the deficiency of the XPC protein^17^. To validate these traits and replicate XP-C disease, we initially examined the photosensitivity profile after UVB exposure in wild-type and XPC knockout keratinocytes, fibroblasts, and melanocytes. Cells from each type were cultured until reaching 80% confluence and then subjected to varying doses of UVB (100, 200, 500, 1000, 4000 J/m2 for keratinocytes and melanocytes, and 200, 300, 500, 700, 1000, 4000, 15000 J/m2 for fibroblasts). This allowed us to assess photosensitivity in a dose- and time-dependent manner (24, 48, and 72 hours). Elevated doses of UVB exposure led to decreased viability in all cell types, including both wild-type and XPC knockout (KO) cells. XPC KO keratinocytes and melanocytes exhibited an increased photosensitivity, displaying a significantly sharper (p<0.001) decline in viability at each UVB dose and across the specified time intervals compared to their wild-type counterparts. Conversely, there was minimal disparity in photosensitivity between wild-type and XPC KO fibroblasts across the three-time intervals (24, 48, and 72 hours).

In the case of keratinocytes, XPC KO cells demonstrated a lower LD50 (154.3 J/m^2^) compared to wild-type cells, which required a substantially higher dose (353.6 J/m^2^) to induce an equivalent 50% lethality after 24 hours of UVB irradiation (Figure 5A). Following 48 hours of UVB exposure, XPC KO cells displayed a more hypersensitive phenotype compared to both wild-type cells and their own state at 24 hours, evident in the divergence of the graphical curves between the two cell types. Additionally, XPC KO cells exhibited a reduced LD50 (102.9 J/m^2^) in contrast to wild-type cells, which necessitated a substantially higher dose (317.2 J/m^2^) to achieve an equivalent 50% lethality after 48 hours of UVB irradiation (Figure 5B). Moreover, following 72 hours of UVB exposure, XPC KO cells demonstrated the most pronounced hypersensitive phenotype compared to wild-type cells, as well as in comparison to the 24- and 48-hour time points. This heightened sensitivity is evident in the separation of the graphical curves between the two cell types, particularly noticeable at a dose of 200 J/m^2^. Furthermore, XPC KO cells displayed a lower LD50 (135.2 J/m^2^) relative to wild-type cells, which required a significantly higher dose (329.5 J/m^2^) to induce an equal 50% lethality after 72 hours of UVB irradiation (Figure 5C).

**Figure 5.**
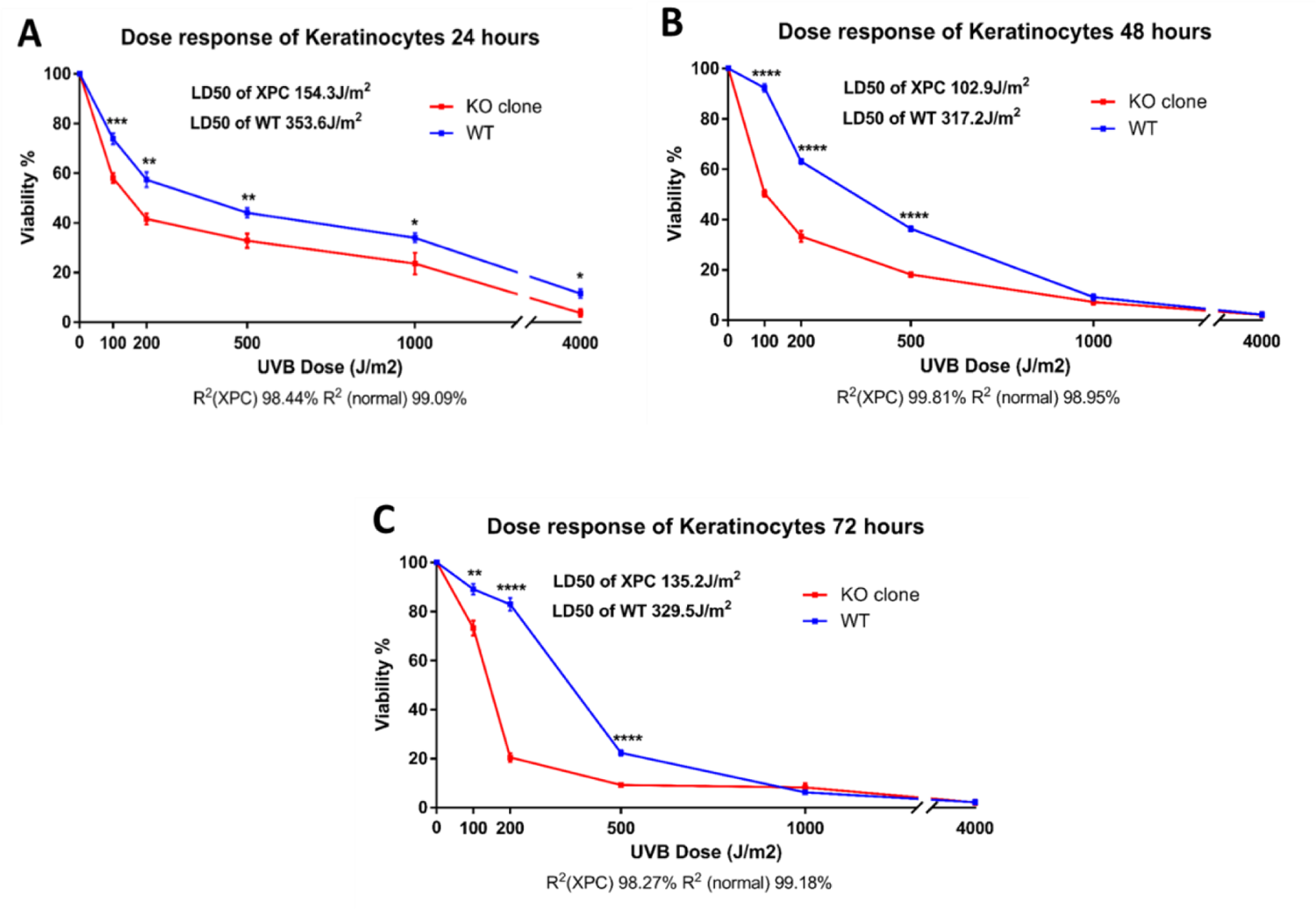
XPC KO N/TERT-2G keratinocyte cells manifest increased hypersensitivity to UVB irradiation in a dose and time dependent manner. (A) Viability of keratinocytes 24h post UVB irradiation. (B) Viability of keratinocytes 48h post UVB irradiation. (C) Viability of keratinocytes 72h post UVB irradiation. Both XPC KO and wild-type cells were seeded in 6 well plates to be irradiated at 80% confluence with increasing UVB doses. Cells were exposed to various doses of UVB and their viability was assessed after 24, 48, and 72 hours through incubation with trypan blue. XPC knockout (KO) cells demonstrated a notably steeper and statistically significant reduction in viability with rising UVB doses compared to wild-type cells. Viability was determined as a percentage of the control, with non-irradiated cells representing 100% viability. Statistical analysis revealed a highly significant difference *p-value<0.05, **p-value<0.01, ***p-value<0.001, ****p-value<0.0001 (unpaired t-test). The reported results are the average of three separate biological experiments (N=3).

In the case of melanocytes, XPC knockout (KO) cells exhibited a reduced LD50 (251.3 J/m^2^) in contrast to wild-type cells, which needed a significantly higher dose (848.6 J/m^2^) to achieve a 50% lethality rate after 24 hours of UVB exposure (Figure 6A). Following 48 hours of UVB exposure, XPC KO cells displayed more photosensitivity compared to wild-type cells, as well as in comparison to their sensitivity at 24 hours. This heightened sensitivity is evident from the distinct separation of the graphical curves between both cell types, especially beyond a dose of 100 J/m^2^. at Furthermore, XPC KO cells also demonstrated a decreased LD50 (79.43 J/m^2^) compared to wild-type cells, which required a significantly higher dose (244.7 J/m^2^) to achieve a 50% lethality rate after 48 hours of UVB exposure (as indicated in Figure 6B). Additionally, after 72 hours of UVB exposure, XPC KO cells displayed the most pronounced hypersensitive phenotype when compared to wild-type cells and their state at 24 and 48 hours. This increased sensitivity is evident from the clear divergence of the graphical curves between both cell types, particularly at a dose of 100 J/m^2^, where the viability of XPC KO cells dropped to nearly 20%, while wild-type cells remained at around 100%. Moreover, XPC KO cells exhibited the lowest LD50 (33.82 J/m^2^) in contrast to wild-type cells, which needed a considerably higher dose (519.9 J/m^2^) to achieve a 50% lethality rate after 72 hours of UVB exposure (Figure 6C).

**Figure 6.**
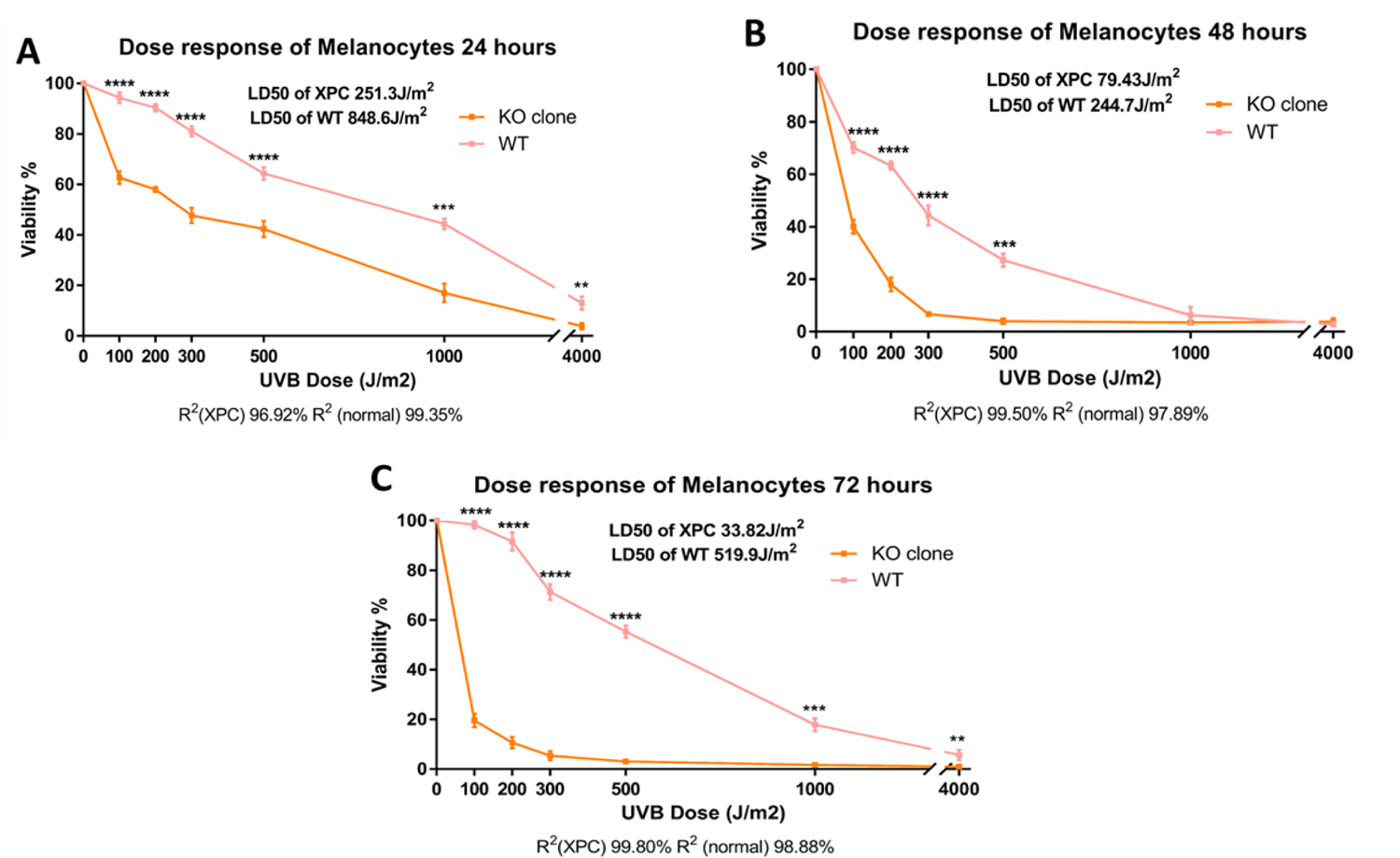
XPC KO Mel-ST melanocyte cells manifest increased hypersensitivity to UVB irradiation in a dose and time dependent manner. (A) Viability of melanocytes 24h post UVB irradiation. (B) Viability of melanocytes 48h post UVB irradiation. (C) Viability of melanocytes 72h post UVB irradiation. Both XPC KO and wild-type cells were seeded in 6 well plates to be irradiated at 80% confluence with increasing UVB doses. Cells were exposed to varying doses and their viability was assessed at 24, 48, and 72-hour intervals using trypan blue incubation. XPC knockout (KO) cells exhibited a more pronounced and statistically significant reduction in viability as the UVB dose increased, in contrast to wild-type cells. Viability was determined by calculating the percentage in relation to the control, where non-irradiated cells represented 100% viability. The statistical significance was denoted as **p-value<0.01, ***p-value<0.001, ****p-value<0.0001 (unpaired t-test). The findings are based on the average of three independent biological replicates (N=3).

In the case of fibroblasts, the photosensitivity profile was somewhat less pronounced compared to what was observed for keratinocytes and melanocytes. Specifically, XPC knockout (KO) cells exhibited a slightly lower LD50 (1359 J/m^2^) in contrast to wild-type cells, which required a slightly higher dose (1778 J/m^2^) to achieve a 50% lethality rate after 24 hours of UVB irradiation (Figure 7A). Following 48 hours of UVB exposure, XPC KO cells displayed a very slight, yet noticeable, hypersensitivity when compared to wild-type cells, especially at a dose of 4000 J/m^2^. Additionally, XPC KO cells also showed a lower LD50 (948.5 J/m^2^) compared to wild-type cells, which required a slightly higher dose (1182 J/m^2^) to reach a 50% fatality rate after 48 hours of UVB irradiation (Figure 7B). Finally, after 72 hours of UVB irradiation, XPC KO cells exhibited a heightened sensitivity compared to both wild-type cells and their sensitivity at 24 and 48 hours. Moreover, XPC KO cells demonstrated the lowest LD50 (655.7 J/m^2^) in contrast to wild-type cells, which needed a considerably higher dose (923.9 J/m^2^) to achieve a 50% fatality rate after 72 hours of UVB irradiation (Figure 7C).

**Figure 7.**
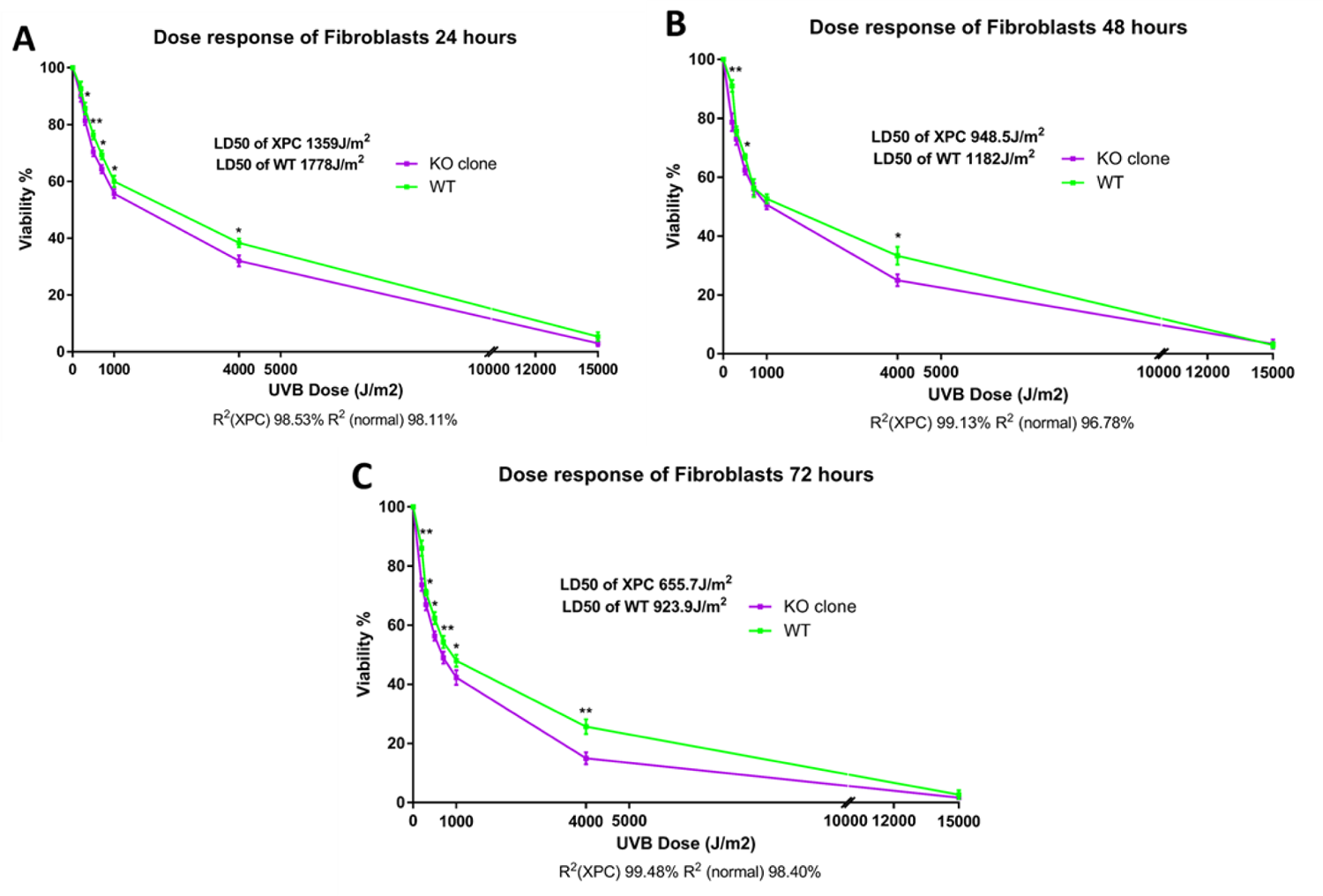
XPC KO S1F/TERT-1 fibroblast cells manifest a slight hypersensitivity to UVB irradiation in a dose and time dependent manner. (A) Viability of fibroblasts 24h post UVB irradiation. (B) Viability of fibroblasts 48h post UVB irradiation. (C) Viability of fibroblasts 72h post UVB irradiation. Both XPC KO and wild-type cells were seeded in 6 well plates to be irradiated at 80% confluence with increasing UVB doses. The cells were exposed to varying doses, and their viability was assessed after 24, 48, and 72 hours by utilizing trypan blue incubation. Notably, XPC knockout (KO) cells exhibited slight statistically significant reduction in viability as the UVB dose increased, as compared to the wild-type cells. Viability was determined by calculating the percentage in relation to the control, where non-irradiated cells were considered to have 100% viability. The statistical significance was indicated as *p-value<0.05, **p-value<0.01 (unpaired t-test). The presented results are the average of three independent biological replicates (N=3).

#### 6-4PPs Repair Capacity

As previously mentioned, XPC protein plays a pivotal role as a key sensor in the initial recognition phase of the global genome repair pathway, which addresses UV-induced photoproducts^17^. Our objective was to investigate how the XPC KO mutation impacted the DNA damage repair kinetics resulting from UVB irradiation in the three types of XPC KO cells (keratinocytes, fibroblasts, and melanocytes) to model XP-C disease. UVB exposure primarily generates two major photo-lesions: 6-4 photoproducts (6-4PPs) and cyclobutane pyrimidine dimers (CPDs). These two lesions exhibit distinct structural characteristics, with 6-4 photoproducts (6-4PPs) causing a more pronounced distortion of the DNA helix (a 44° bend) compared to cyclobutane pyrimidine dimers (CPDs), which induce a milder helix distortion of (a 9° bend). The differential distortion in DNA makes it easier for XPC to recognize 6-4PPs compared to CPDs. Furthermore, 6-4PPs are considerably more efficiently repaired by nucleotide excision repair (NER) mechanisms, with a half-life of 2 hours for 6-4PPs compared to 33 hours for CPDs^4^. Therefore, for future experiments, we determined that focusing on 6-4PPs would be a more compelling approach for assessing the outcomes and can be of main interest for future therapeutic strategies taking advantage of the short follow-up timing accompanied with the 24 hour readout following UVB exposure as shown here^23^, as CPDs require a longer follow-up time.

To assess DNA damage, wild-type and XPC KO N/TERT-2G keratinocytes and Mel-ST melanocytes were exposed to a dose of 150 J/m^2^, while S1F/TERT-1 fibroblasts were exposed to a dose of 300 J/m^2^. This choice of doses was based on their relatively higher resistance compared to melanocytes and keratinocytes, as determined through dose-response curve analysis. The purpose of using these doses was to induce DNA damage adaptable with the percentage of cellular mortality, allowing for the quantification of 6-4PPs through single-cell fluorescence analysis, measuring the relative fluorescence unit (RFU) after immunofluorescence staining.

At the initial 0-hour time point after exposure to UVB irradiation, immunocytochemistry staining of 6-4PPs revealed comparable levels of accumulated DNA damage in XPC KO keratinocytes (Figure 8A), fibroblasts (Figure 8B), and melanocytes (Figure 8C). However, after 24 hours of UVB irradiation, wild-type cells demonstrated the ability to repair 6-4PPs, unlike XPC KO cells, which exhibited a deficiency in 6-4PPs repair. This indicates a noticeable lag or absence in the 6-4PPs repair capacity of XPC KO cells compared to their corresponding wild-type cells.

**Figure 8.**
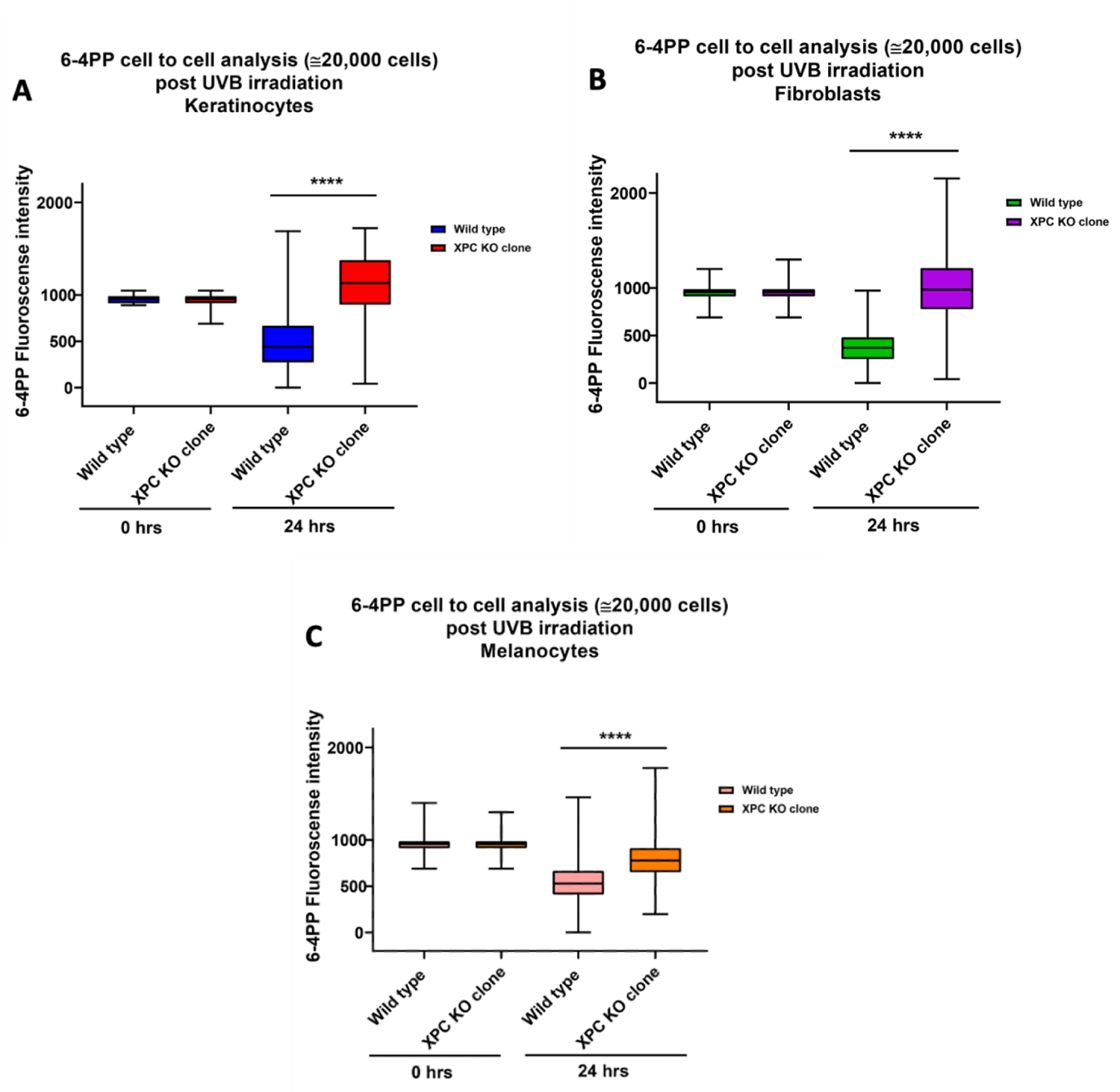
XPC KO N/TERT-2G keratinocyte, S1F/TERT-1 fibroblast, and Mel-ST melanocyte cells manifest a significantly persistent and unrepaired 6-4PPs post 24 hours of UVB irradiation. (A,B,C) 6-4PPs repair assay in wild-type and XPC KO keratinocytes, fibroblasts, and melanocytes. XPC knockout in keratinocytes (A), fibroblasts (B), and melanocytes (C) resulted in a notably persistent and unrepaired UVB-induced 6-4PPs lesions (fluorescence intensity) compared to their respective wild-type cells 24 hours after UVB irradiation. It is noteworthy that all cell types displayed similar levels of 6-4PPs at 0 hours post-UVB irradiation. To test the repair capacity, both wild type and XPC KO cells from each cell line and type were seeded to reach 80% confluence. Afterwards, these cells were subjected to UVB irradiation. Following UVB irradiation at time 0h and after 24 hours, cells were further stained based on the protocol, which comprises the fixation of the cells using 4% paraformaldehyde and 0.2% of Triton X-100 to permeabilize the cells. 2M HCL was then utilized to fully denature the DNA double helix, enhancing the access of the antibody targeting DNA damage caused by UVB irradiation. After the saturation process, cells were incubated overnight with primary 6-4PP antibody. Secondary mouse antibody FITC was then added the next day. Single-cell analysis was carried out via quantifying nuclear DNA damage in several individual cells per condition and were constructed as box plots. **** p<0.0001 unpaired t-test. The reported results are the average of three separate biological experiments (N=3).

#### Proliferation Status

After a brief period following XPC knockout, our microscopic analysis revealed a visually significant halt in the proliferation/growth capacity of the three types of XPC knockout skin cells, as opposed to their respective wild-type counterparts. To confirm this impact, we conducted a 5-hour EDU assay on both wild-type and XPC KO cell types, followed by the analysis of EDU incorporation. It was observed that wild-type cells exhibited significantly higher EDU incorporation, implying more proliferation in keratinocytes (Figure 9A), fibroblasts (Figure 9B), and melanocytes (Figure 9C) compared to the knockout clones. Out of 100% for keratinocytes, wild-type EDU-positive cells were 51%, whilst 32% for the XPC KO cells. Out of 100% for fibroblasts, wild-type EDU-positive cells were 55%, whilst 41% for the XPC KO cells. Out of 100% for melanocytes, wild-type EDU-positive cells were 64%, whilst 44% for the XPC KO cells. This emphasizes the significance of XPC in influencing the multiplication of skin cells, presenting it as a novel strategy for characterization as well.

**Figure 9.**
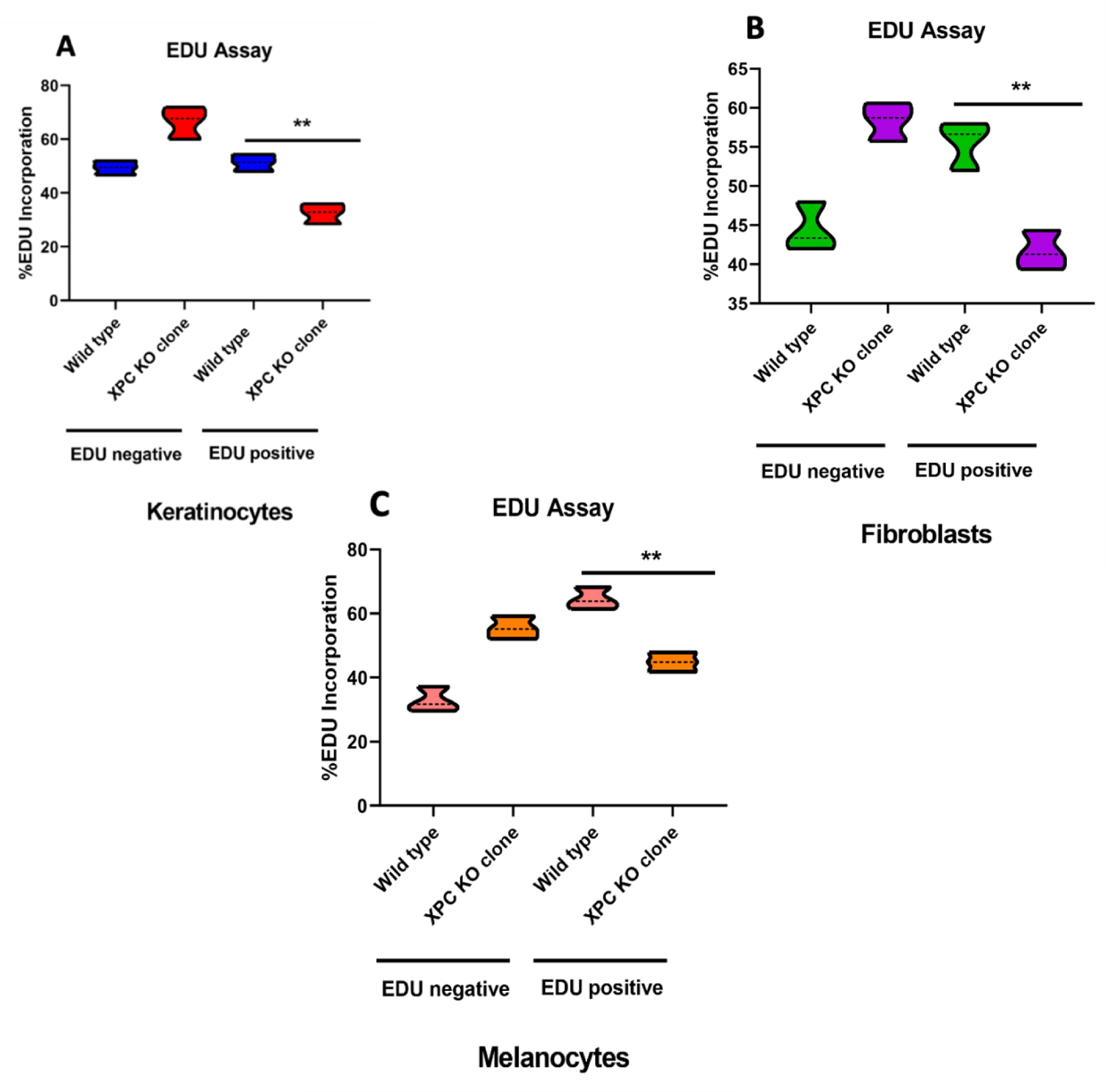
XPC KO manifests a partial halting in the proliferative capacity of N/TERT-2G keratinocytes, S1F/TERT-1 fibroblasts, and Mel-ST melanocytes. (A,B,C) EDU incorporation assay for wild-type and XPC KO keratinocytes, fibroblasts, and melanocytes. An EDU assay was carried out to determine the effect of XPC KO mutation on the proliferative capacity of human immortalized skin cells (keratinocytes, fibroblasts and melanocytes). The incorporation of the nucleoside analog EDU into the cells can be used to determine the health and genotoxicity of the cells. A click-it covalent reaction between an azide and an alkyne, which copper catalyzes, can further quantify the incorporation of the nucleoside analog EDU. To do that, wild-type and XPC KO of each cell type were seeded in 6 well plates to reach 50% of confluence. Afterwards, EDU was diluted and added to the cell media for 5 hours. Cells were then trypsinized, harvested, and stained according to the manufacturer’s protocol. The readout was done using flow cytometry (FACScan, BD LSRII flow cytometer, BD Biosciences). The post-analysis was done using flowing software (Turku Bioimaging, Finland). **p-value<0.01 (unpaired t-test). The results presented are the mean of three biological replicates (N=3).

### XPC KO Induces ECM Scaffold Degradation in 3D Reconstructed Skin Models

Conducting research on cells in a 2D culture is essential, but this approach is limited by its inability to accurately replicate conditions found in the human body. Additionally, the rarity of XP-C disease poses challenges in obtaining skin samples from patients for analysis. As an alternative, a 3D reconstructed skin model with XPC knockout (KO) cells derived from amplified 2D cultures can be generated. This model offers researchers a more physiologically relevant representation and provides the opportunity to employ additional constructs for experimentation, enabling a deeper understanding of the mechanisms activated by XPC mutation. The workflow (Figure 10) consists of seeding separately wild-type and XPC KO fibroblasts, being embedded in a specific gel termed fibrin, in an insert support to permit the formation of the dermal equivalent. This process lasts 10-12 days, allowing sufficient proliferation and extracellular matrix production. Afterwards, wild-type and XPC KO melanocytes and keratinocytes are mixed together (each separately), and both cell types are seeded on the top of either the wild-type or XPC KO dermal equivalent and kept for 4-5 days to proliferate in immerged culture. After that, they are switched to the air liquid interface stage for 10 days or more, where media is aspirated, allowing the epidermal differentiation process to produce either a full wild type or XPC KO 3D reconstructed skin model. At the step of the differentiation process, we observed that all the fibrin gels (N=10) of XPC KO skin cell types started to extensively degrade (Figure 10) compared to the wild type which did not show any of these features and had succeeded to harvest the skin architecture (data not shown here).

**Figure 10.**
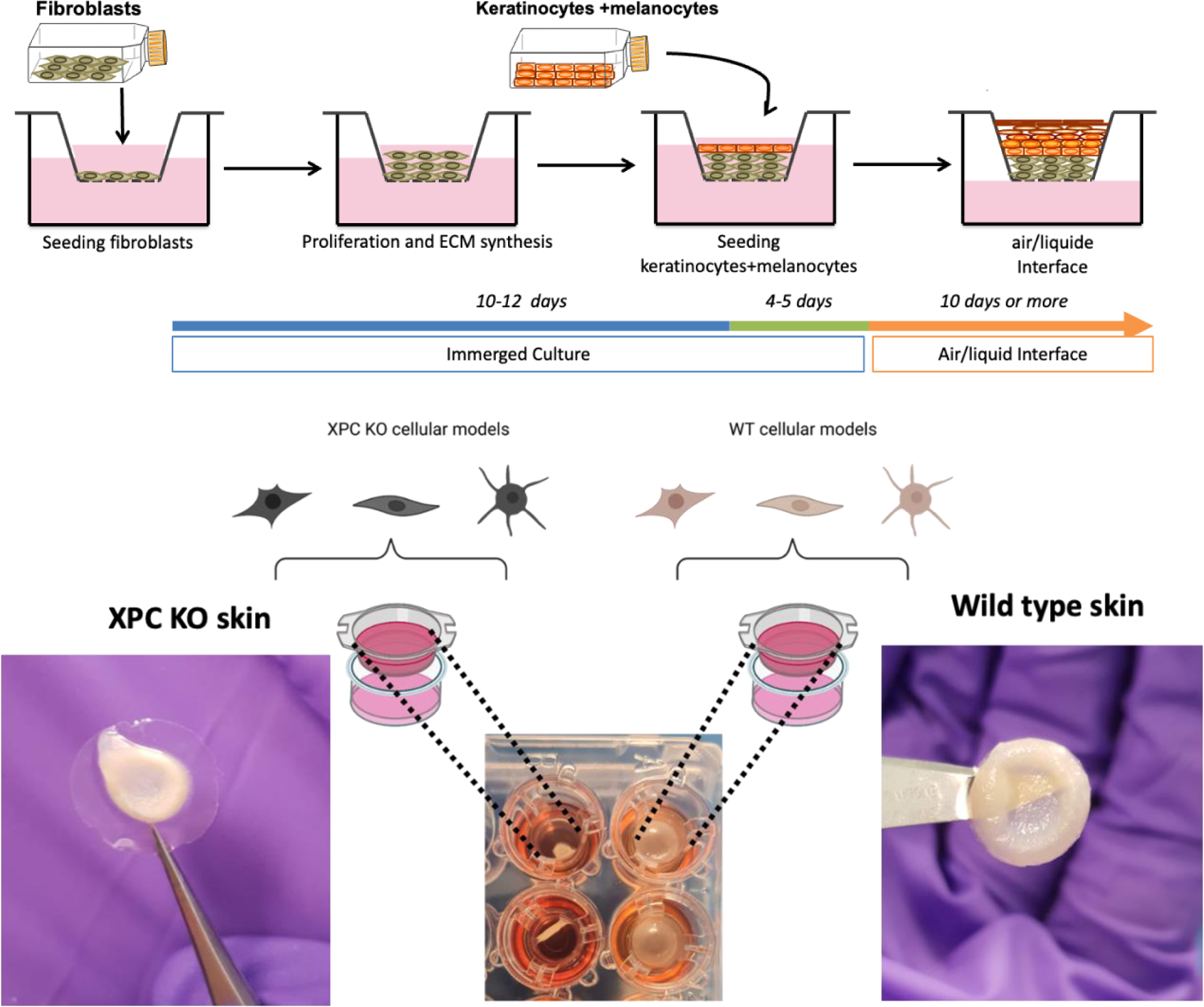
Degradation of the fibrin gel in XPC KO skin model during differentiation processes. The protocol consists of seeding separately wild type and XPC KO fibroblasts, being embedded in a specific gel termed fibrin, in insert support to permit the formation of the dermal equivalent. This process lasts 10-12 days, allowing sufficient proliferation and extracellular matrix production. Afterwards, wild-type and XPC KO melanocytes and keratinocytes are mixed (each separately), and both cell types are seeded on the top of either the wild-type or XPC KO dermal equivalent and kept for 4-5 days to proliferate in immerged culture. After that, they are switched to the air-liquid interface stage for 10 days or more, where media is aspirated, allowing the epidermal differentiation process. During the last step, an extensive degradation and retraction of XPC KO scaffold was observed. The results presented are the mean of three biological replicates (N=10).

### XPC KO Induces an Inflammatory Secretome Profile triggered by Fibroblasts

To formulate an initial interpretation of our observations, we referred to the findings reported by Thierry Magnaldo’s group which indicated an increase in the expression of matrix metalloproteinase 1 (MMP-1) in XP-C fibroblasts^24^. To extend our analysis, we aimed to analyze a bunch of inflammatory cytokines (IL-1β, IFN-α2, IFN-γ, TNF-α, MCP-1, IL-6, IL-8, IL-10, IL-12, IL-17A, IL-18, IL-23, and IL-33) from the secretome, being well known to be stimulators of MMPs^25,26^ of our XPC KO fibroblasts versus their associated wild-type cells following 24 hours of culture. Strikingly, we observed that almost all these cytokines were strongly induced following XPC KO implementing a strong inflammatory shift (Figure 11). In XPC knockout fibroblasts, the concentration of IFN-α2 experienced a significant increase to 13,566 pg/ml, marking a 1.5-fold rise compared to their respective wild-type counterparts, which had a concentration of 8,982 pg/ml. Similarly, for IFN-γ, the concentration surged to 8,866 pg/ml, indicating a 1.5-fold increase compared to the corresponding wild-type level of 5,935 pg/ml. As for IL-1β, its concentration dramatically increased to 11,000 pg/ml, showcasing a substantial 4.64-fold rise compared to the wild-type concentration of 2,368 pg/ml. IL-6 exhibited a noteworthy increase to 29 pg/ml, representing a substantial 6.50-fold elevation compared to the 6 pg/ml concentration in the wild-type counterparts. The concentration of IL-8 rose significantly to 43 pg/ml, reflecting a 5.3-fold increase compared to the 8 pg/ml concentration in their wild-type counterparts.

**Figure 11.**
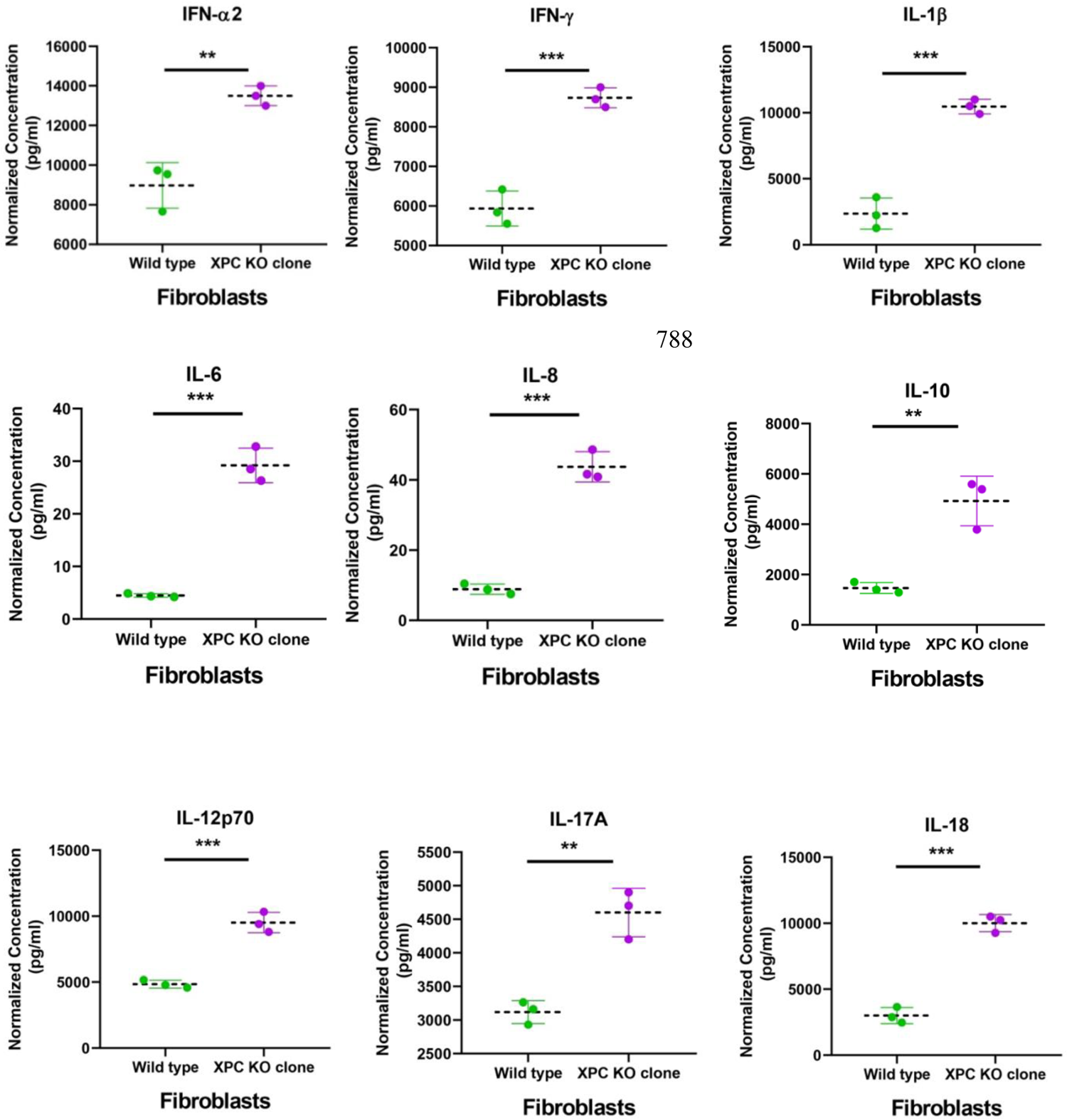
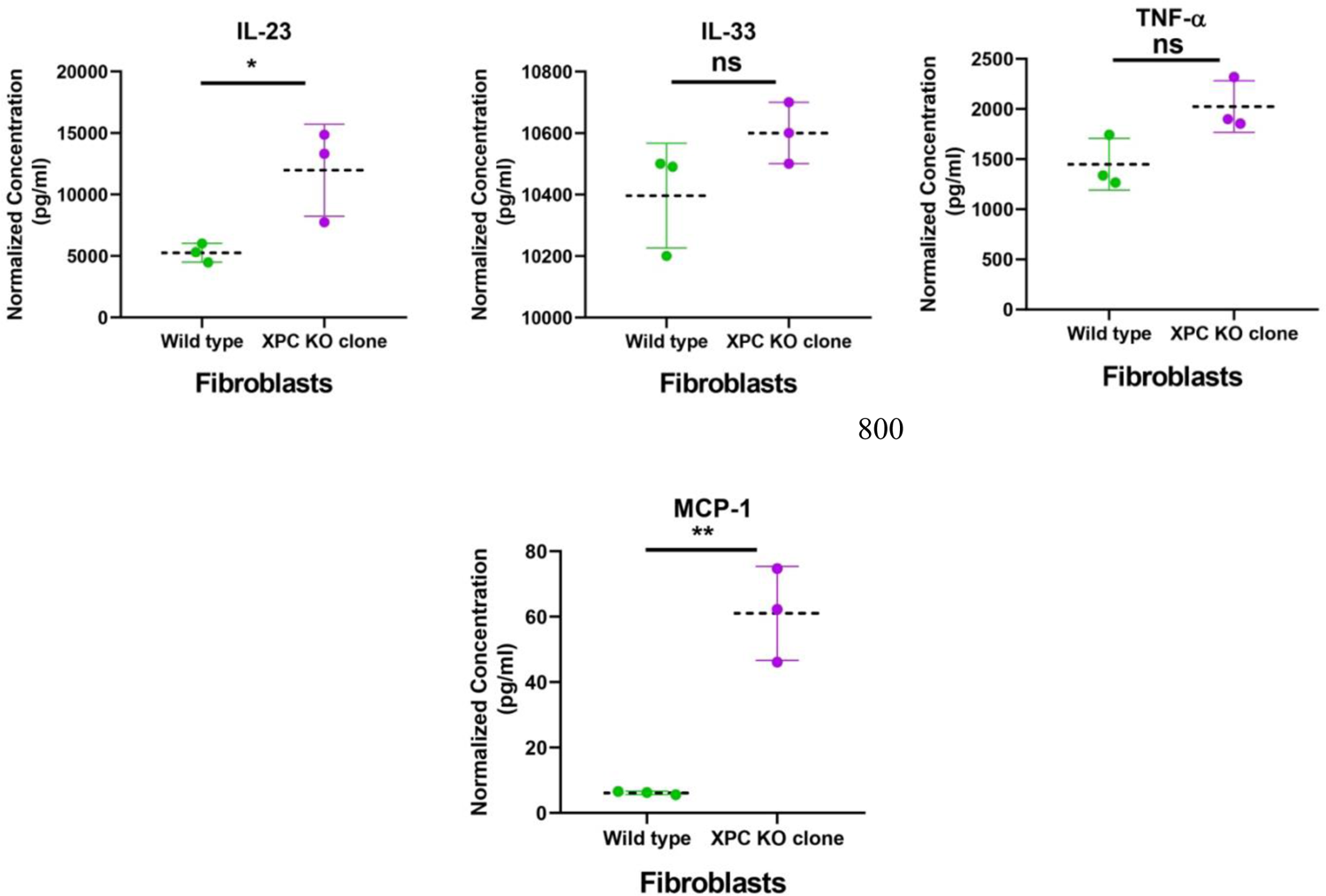
XPC KO induces a rise in the inflammatory secretome signature secreted by S1F/TERT-1 fibroblasts. XPC KO in S1F/TERT-1 fibroblasts induce an increase in the human inflammatory secretion profile encompassing both chemokines and cytokines. This included a rise in IL-1β, IFN-α2, IFN-γ, TNF, MCP-1 (CCL2), IL-6, IL-8 (CXCL8), IL-10, IL-12p70, IL-17A, IL-18, IL-23, and IL-33. Subsequently, samples were acquired using a FACSCanto™ II cytometer and analyzed utilizing the online QOGNIT LEGENDplex™ program. The statistical significance was denoted as **p-value<0.01, ***p-value<0.001, ****p-value<0.0001 (unpaired t-test). The results presented are the mean of three biological replicates (N=3).

Moreover, IL-10 showed an increased concentration of 4,922 pg/ml, marking a 3.3-fold rise compared to the corresponding wild-type concentration of 1,464 pg/ml. IL-12p70 demonstrated a significant increase to 9,511 pg/ml, indicating a 2-fold rise in comparison to the wild-type concentration of 4,845 pg/ml. Similarly, IL-17A exhibited a concentration increase to 4,600 pg/ml, representing a 1.5-fold elevation compared to the wild-type concentration of 3,118 pg/ml. IL-18 displayed a substantial increase to 10,000 pg/ml, showcasing a notable 3.33-fold rise compared to the wild-type concentration of 2,999 pg/ml. IL-23 exhibited a significant increase to 11,970 pg/ml, reflecting a 2.3-fold elevation compared to the 5,262 pg/ml concentration in their wild-type counterparts. In contrast, IL-33 showed no significant difference in concentration, hovering around 10,500 pg/ml. While TNF-α demonstrated a slight increase in concentration to 1,882 pg/ml, it did not reach statistical significance, representing a 1.3-fold rise compared to the wild-type concentration of 1,448 pg/ml. Finally, MCP-1 displayed a substantial increase to 61 pg/ml, indicating a remarkable 9.95-fold rise compared to the wild-type concentration of 6 pg/ml.

## Discussion

Examining the function of a gene or multiple genes in human cells is essential for understanding the complex mechanisms behind various human diseases. Many research studies have utilized RNA interference (RNAi) technology to diminish the expression of specific genes^27,28^. However, this approach, which primarily aims at knockdown, still needs to fully attain a total elimination of gene/protein expression and may result in unintended off-target effects^29^. Consequently, there is a pressing need for techniques that enable the comprehensive knockout of a gene in human cells. Our research work marked for the first time a simple, straightforward, and successful modeling strategy for the XP-C disease in all three immortalized cell types which form the main core of the human skin—keratinocytes, fibroblasts, and melanocytes. This was achieved by introducing a loss-of-function mutation to the *XPC* gene at the exon 3 site through the CRISPR-Cas9 strategy based on ribonucleoprotein (RNP), utilizing the same single-guide RNA (sgRNA) via NEON electroporation system. For maximal delivery and editing efficacy, three parameters must be roughly optimized, and each parameter needs to complement the other. These include delivery approach, cargo system, and cellular type (proliferation rate, fragility, and size). Based on the literature, the ribonucleoprotein (RNP) approach coupled with NEON electroporation exerts a strong editing potential^30^. RNP complex is simple to create, has a transient transfection impact that doesn’t integrate into the host’s DNA, limiting the off-target effect, and exerts minimal cytotoxicity thanks to its small size contrary to that of the plasmid strategy^31^. NEON electroporation is categorized as a physical delivery approach that can be optimized to select the best parameters (number of pulses, time, and voltage) suitable for the maximal delivery efficacy to the host cells of interest, giving it a strong advantage over the other electroporation systems like nucleofector™ II/2b device that lacks the setup parameters option, thus can drastically cause enormous cellular stress and mortality. For our manipulated human skin cells (keratinocytes, fibroblasts, and melanocytes), several NEON programs were tested. Finally, 1700v, 10ms, 1pulse was the best, along with the RNP system, to perturb the *XPC* gene reaching an editing efficacy of around 99%. This efficacy yield facilitated the selection of the homogenous knockout clones from the three heterogenous (edited and non-edited) cell types. It’s important to note that this editing strategy can be applied to pursue different research objectives. By adjusting the specific sgRNA of interest, researchers can use various cell types and target different genes.

XP-C stands out as one of the prevalent types of XP^32^. Almost all of these mutations can be in the form of nonsense, frameshift, and deletion events in XP-C patients^33^. In XP-C patients, there is an absence of XPC protein and almost a negligible expression of XPC at the mRNA level^33^. Our knockout results were confirmed for the three cell types at the protein level using western blot assay, where they showed a complete absence of the XPC protein band compared to their associated wild-type controls. This result is consistent with the prior study conducted by Chavanne et al.^34^, which reported the absence of XPC protein in cells obtained from individuals with XP-C. Furthermore, the significant downregulation of the XPC mRNA expression level was also confirmed via RT-PCR for the three cell types. Legerski and Peterson^35^, Khan et al.^33^, and Fayyad et al.^18^ reported minimal levels of XPC mRNA in cells derived from XP-C patients. This indicates that CRISPR-Cas9 mediated edit of *XPC* gene can tend to generate mutant mRNAs that may harbor premature termination codons (PTCs) that trigger nonsense-mediated mRNA decay (NMD pathway) as a protective mechanism, preventing the expression of harmful truncated proteins. This suggestion has been also implemented by several groups that worked on cells from XP-C patients^18,33,34,36^. Using immunofluorescence staining, we confirmed the presence of XPC protein in the nucleus for the wild-type cells; this is not surprising since this protein is a critical player in the GG-NER system^37^, and its complete absence following the knockout in the three cell types. Finally, Sanger sequencing showed that the exon 3 region of the *XPC* gene harbors 2 nucleotides indel mutation for keratinocytes, 5 nucleotides indel mutation for fibroblasts, and 258 nucleotides indel mutation for melanocytes (data not shown here). These indel mutations disrupted the open reading frame by yielding an early stop codon. A two-stage mechanism can account for the homozygous mutation. Because it is unlikely that the homozygous indel mutation was caused by two simultaneous deletion events in both alleles, we hypothesized that it was caused by a loss of one allele or by a sequential process of an initial deletion in one allele caused by non-homologous end joining (NHEJ) followed by repair of the other allele by the homology-directed repair (HDR) using the altered allele sequence as a template.

Research articles investigating the link between XPC, and skin biology concentrate on primary XPC-mutated keratinocytes and fibroblasts^17,18^. As previously highlighted, the absence of an appropriate control in primary XP-C cells hinders the comparison of outcome data, posing a challenge in exploring the molecular events associated with this disease, especially given the inherent heterogeneity among individuals and we should be aware that since the GG-NER DNA repair system is absent in XP-C patients, a higher susceptibility to other unknown genetic mutations can occur hindering the precise outcomes^7^. Therefore, the utilization of CRISPR-Cas9 would be advantageous, providing a significant benefit by generating a mirrored control and facilitating precise outcome results that are specifically related to the loss of XPC. Unfortunately, due to the challenging manipulation of primary XPC-mutated melanocytes within their limited passage potential, there is currently no available data in the literature that highlights their response to XPC mutation. Given that Xeroderma Pigmentosum C disease is characterized by abnormalities in pigmentation in patients, the necessity for a diseased melanocyte model becomes evident to explore the molecular pigmentation features associated with this disease since the term Pigmentosum implies abnormalities in pigmentation present in patients^38^.

Here, the XPC knockout skin cell types, encompassing keratinocytes, fibroblasts, and melanocytes, underwent characterization across three levels: UVB photosensitivity, 6-4PPs repair capacity, and proliferation status. What was of main interest was the distinct behavioral responses, particularly the variation in photosensitivity intervals, among these three cell types when exposed to UVB irradiation. In our study on N/TERT-2G keratinocytes, we compared the viability of XPC KO with their associated wild-type cells under a time (24, 48, and 72 hours) and UVB dose dependent manner, leading us to confirm their photosensitive nature. Additionally, our analysis of UVB-induced DNA lesions (6-4PPs) through single-cell quantification using immunofluorescence showed that XPC KO keratinocytes experienced a disturbance in their GG-NER system compared to their associated wild-type cells 24 hours after UVB irradiation. This perturbation was evident as the lesions persisted, indicating impaired DNA repair. Remarkably, our findings align closely with those of Warrick et al.^17^, who studied three independent XP-C mutated keratinocytes from patients and characterized this model. This concurrence in results further supports the validity of our XPC KO keratinocyte model and its resemblance to the real XP-C disease phenotype. Given that keratinocytes are crucial in the pathology of XP-C disease, being the uppermost cells exposed to UV radiation and capable of transforming into non-melanoma skin cancers, our CRISPR-Cas9 system-generated model can provide valuable insights into the specific disrupted pathways that are uniquely altered in XPC KO keratinocytes compared to wild-type cells. These insights can potentially advance our understanding of XP-C disease and may contribute to the development of novel therapeutic approaches as shown in our recent work by Kobaisi et al.^23^ In the case of S1F/TERT-1 fibroblasts, we observed a very slight/nearly absence but significant difference in viability between wild-type and XPC KO fibroblasts. Interestingly, both wild-type and XPC KO fibroblasts exhibited a similar pattern of decreasing viability, in response to increasing UVB exposure in a time (24, 48, and 72 hours) and dose-dependent manner, highlighting a nearly slight to absence of photosensitive nature when XPC is lost. Furthermore, our single cell quantification analysis of UVB-induced DNA lesions (6-4PPs) using immunofluorescence after 24 hours of UVB irradiation revealed that XPC KO fibroblasts exhibited a perturbation in their GG-NER system compared to their associated wild-type cells like what was observed in the study by Fayyad et al. In Fayyad’s work, three independent primary XP-C mutated fibroblasts from patients were characterized, and their responses to UVB dose-dependent photosensitivity were almost identical to those of the wild-type fibroblasts, showing no significant difference^18^. Our immortalized XPC KO fibroblast model demonstrated strong coherency with their findings, supporting the validity of our approach to mimic the real disease phenotype from patients. De Waard et al.^39^ also demonstrated similar findings as Fayyad et al. regarding photosensitivity in their study. They showed that XP-C mutated fibroblasts exhibited comparable photosensitivity to wild-type fibroblasts with an accumulation of 6-4PPs damage following 24 hours of UVB irradiation. XPC KO fibroblasts generation can be of main interest given the fact that these cells are well known to influence keratinocytes and melanocytes through their secretions and might serve a more realistic model following coculturing or 3D experimental designs linked to study XP-C disease and follow-up of skin cancer onset. In our study on melanocytes, we observed the most pronounced photosensitivity profile in XPC KO melanocytes compared to wild-type cells, demonstrating a clear response in a time (24, 48, and 72 hours) and dose-dependent manner. Additionally, our examination of UVB-induced DNA lesions (6-4PPs) through single-cell quantification analysis, conducted using immunofluorescence after 24 hours of UVB exposure, demonstrated that melanocytes lacking XPC exhibited a disturbance in their GG-NER system when compared to their corresponding wild-type cells. It’s important to note that the lack of data in the literature on XP-C mutated melanocytes from patients necessitated us to develop our own protocol for modeling this disease in melanocytes. XPC KO melanocytes model can be of main interest for future profiling of the pigmentation machinery to develop photoprotective strategies linked to XP-C disease and why not to various pigmentary disorders since this domain is still elusive. Indeed, for the proliferation part, our microscopic observation urged an EDU assay to prove the effect of the XPC KO on partially halting the proliferation of XPC KO keratinocytes, fibroblasts, and melanocytes. Here, we show a new possible method for characterization of XPC mutants via assessing the effect on the proliferation status specifically in the skin biology context. Indeed, some papers have studied the impact of XPC silencing on the proliferation status of lung cancerous cells^40,41^. Cui et al. have shown that XPC silencing in non-small-cell lung cancer (NSCLC) cells has increased their proliferation and migration status ^40^ while Teng et al. have shown that XPC downregulation will tend to promote more sensitivity towards cisplatin treatment, implying a decreased proliferation of adenocarcinoma cells^41^. Due to its multifunctional nature, XPC may demonstrate diverse effects depending on the organ, requiring a specific emphasis on its association with proliferation in each compartment individually to understand the overall impact. In this study, we highlight for the first time that the knockout of XPC disrupts the proliferation of human skin cells including keratinocytes, fibroblasts, and melanocytes.

3D disease modeling offers more realistic environment to study interactions^42^. Constructing a skin model with immortalized human cells faces its greatest challenge in achieving the various epidermal layers and by preserving a proper differentiation process. For instance, HaCaT cells have proven to encounter difficulties in replicating primary human skin cells^43^ due to it’s chromosomal abnormalities, whereas the N/TERT-2G cell line has successfully exhibited a substantial similarity to primary human keratinocytes, providing a significant advantage for unrestricted 3D modeling^44^ and demonstrating a normal chromosomal arrangement profile^44^. In a first attempt using XPC KO skin cells, we aimed to generate 3D reconstructed skin model and compare it to its referred wild-type skin model. Unfortunately, a huge degradation of the Extracellular matrix embed in fibrin gel scaffold was observed during the differentiation process compared to the wild-type which demonstrated a well differentiated skin profile (data not shown here). Following that, our attention shifted to XPC KO fibroblasts, being the key cells involved in secretions. Given that Thierry Magnaldo and his team have demonstrated a heightened expression of matrix metalloproteinase 1 (MMP-1) induced by primary XP-C mutated fibroblasts from patients^24^, our goal was to have an explanation. Surprisingly, we observed a strong drive in a bunch inflammatory cytokine (IL-1β, IFN-α2, IFN-γ, TNF-α, MCP-1, IL-6, IL-8, IL-10, IL-12, IL-17A, IL-18, IL-23, and IL-33) present in the secretome of XPC KO fibroblasts versus their associated wild-type cells which might explain this aberrant degradation. IL-1β is a proinflammatory cytokine and has shown to stimulate the production of MMP-1^45,46^. Indeed, Sánchez et al. has also shown that increased production of both IL-1β and IL-8 was associated with an augmentation in MMPs including MMP-1^47^. Li et al. has shown that IL-6 which is a proinflammatory cytokine can stimulate MMP-1^48^. Du et al. have also proved that TNF-α can stimulate MMP-1 through IL-6 which mediates the effect in fibroblasts^26^. Additionally, Miao et al. have shown that IL-12p70 mediate the expression of MMP-1 via stimulating NF-κB activation pathway^49^. Cortez et al. have also shown that IL-17 induces MMP-1 production in primary human cardiac fibroblasts through the activation of p38 MAPK and ERK1/2, leading to the activation of C/EBP-beta, NF-κB, and AP-1^50^. Wang et al. also demonstrated that IL-18 enhances the release of MMP-1 in human periodontal ligament fibroblasts through the activation of NF-κB signaling^51^. Yamamoto et al. has shown that Monocyte chemoattractant protein-1 (MCP-1) amplifies the gene expression and production of MMP-1 in human fibroblasts through an autocrine loop involving IL-1 alpha^52^. IFN-α2 and IFN-γ are cytokines that are well known to stimulate various inflammatory pathways especially JAK/STAT signaling^53^ that promotes MMP-1 expression. These cytokines have the ability to engage in various combinations of communication and can bind to several receptors associated with inflammatory responses. This could lead us to a preliminary understanding of the anomalous degradation of the scaffold following the knockout of XPC.

## Conclusion

XP-C is a rare inherited genetic disorder characterized by hypersensitivity to ultraviolet radiation and the accumulation of DNA damage. It results from mutations in the *XPC* gene, leading to a deficiency in this protein responsible for recognizing and initiating the repair of UV DNA damage. Children affected by XP-C disease are commonly known as “children of the moon” because they must avoid sunlight to prevent severe sunburns and reduce the significantly increased risk of developing skin cancers, including non-melanoma and melanoma. Unfortunately, there is currently no effective cure for XP-C disease, and management primarily focuses on strict sun protection measures and regular skin screenings to detect cancer early. The journey of a thousand miles begins with one step, here, we demonstrate the building block for a reproducible model for this disease, which can open future avenues for the loss of XPC in the context of skin biology and will permit novel molecular profiling of the mysteries underlying this disease for novel therapeutics and can be used as a tracker for skin cancer onset.

## Materials and Methods

### Cell lines

Human immortalized male epidermal keratinocyte (N/TERT-2G)^54^ and human immortalized male dermal fibroblast (S1F/TERT-1)^54^ cell lines were kindly supplied as a gift from Dr. James Rheinwald Laboratory (Harvard Medical School, Boston, USA). Human immortalized male melanocyte cell line (Mel-ST) was kindly supplied as a gift from Dr. Robert Weinberg Laboratory (Whitehead Institute for Biomedical Research, Cambridge, USA)^55^. Wild type N/TERT-2G cell line was cultured using EpiLife medium with 60 µM calcium (Gibco™, cat. #MEPI500CA), supplemented with human keratinocyte growth supplement (HKGS, Gibco™, cat. #S0015), containing Human growth factor I insulin-like recombinant: 0.01 µg/ml, Bovine pituitary extract (BPE): 0.2% v/v, Bovine transferrin: 5 µg/ml, Hydrocortisone: 0.18 µg/ml and Human epidermal growth factor: 0.2 ng/ml with added CaCl2 (340 µM, Sigma-Aldrich, Saint Louis, USA) and 1% penicillin/streptomycin. Wild type S1F/TERT-1 cell line was cultured using M199 (Gibco™, cat. #11150-059) and M106 (Gibco™, cat. #M-106-500) +15% iron-supplemented newborn bovine calf serum (Hyclone/Thermo Scientific, cat. #SH3007203) +10 ng/ml EGF +0.4 μg/ml hydrocortisone with added 1% penicillin/streptomycin. Wild type Mel-ST cell line was cultured in DMEM (Gibco™, cat. #12430054) supplemented 10% FBS and 1% penicillin/streptomycin. All cell lines were maintained at 37° C in a 5% CO2 incubator. When cells attained confluence, they were passaged 1:4–1:10, depending on the cells utilized. Cells were washed with 8 mL phosphate buffered saline (PBS, pH 7.4, Gibco™) before being dissociated from the culture flask (T75 cm2) with 3 mL 0.05% trypsin/EDTA (Gibco™) for 5-10 minutes at 37°C depending on the cell line. Trypsinization was blocked by the addition of 8 mL of complete culture medium. Furthermore, cells were centrifuged at 100× g speed for 5 minutes and the supernatant was discarded.

### NEON™ ELECTROPORATION SYSTEM

Wild-type N/TERT-2G, S1F/TERT-1 and Mel-ST cell lines were electroporated with ribonucleoprotein complex via Neon™ Electroporation System (Thermofisher Scientific, Massachusetts, USA). To maximize the genome-editing efficacy, 1.5 µg of TrueCut Cas9 protein (Thermofisher Scientific, Massachusetts, USA) was added to 5 μl of resuspension buffer (R buffer), followed by the addition of 300 ng of predesigned sgRNA targeting exon 3 region of the *XPC* gene with a crRNA sequence (5’AGGCACACCATCTGAAGAGA3’) (Thermofisher Scientific, Massachusetts, USA). The sgRNA and Cas9 mixture was incubated for 10 minutes at room temperature to assemble and form the ribonucleoprotein complex. Meanwhile, 100,000 cells were also suspended in 5 μl of R buffer. Then cell suspension was added to the RNP complex mix. 10 μl volume of the mix was then electroporated at 1700v, 10ms, 1pulse, transferred into their pre-warmed media, and incubated for 48 hours in 5% CO2 to expand for the post-genome editing analysis.

### Single-cell limiting dilution

For the selection of single clones, the N/TERT-2G heterogeneous cell population was separated using the standard limiting serial dilution method in a 96-well plate (Greiner Bio-One, France), S1F/TERT-1 and Mel-ST heterogeneous cell populations were sorted using BD FACSMelody™ Cell Sorter. Single-cell clones were marked, tracked and further kept cultured for 2 weeks. Afterwards, single-cell populations were expanded in a 6-well plate (Greiner Bio-One, France) for further post-genome editing analysis.

### RT-qPCR

The total RNA content was extracted from wild-type and XPC KO keratinocytes, fibroblasts and melanocytes using RNeasy plus mini kit (#Cat. 74134, Qiagen, France). Quantification of RNA was done using Nanodrop 1000. Reverse transcription to cDNA was achieved by using RNA (1µg) and the Superscript vilo cDNA synthesis kit (#Cat. 11754050, Invitrogen, Massachusetts, USA). 25 ng of XPC and GAPDH (Qiagen, France) cDNAs were then used to launch the qPCR reaction using gene-specific primers along with the Platinum SYBR green qPCR SuperMix-UDG (#Cat.11733038, Invitrogen, Massachusetts, USA). Using BioRad CFX96TM Real-time System (C1000 TouchTM Thermal Cycler), samples were launched in triplicates (N=3). To unravel the specificity of the primers utilized, Melt curve analysis was conducted to ensure that a single melt-curve peak was present. Glyceraldehyde-3-phosphate dehydrogenase (GAPDH) housekeeping gene was used to normalize the expression level of the target gene. Fold change expression levels were calculated based on the 2-ΔΔCT Livak method. Samples were launched in triplicates (N=3).

### Immunoblotting

Total proteins from wild-type and XPC KO keratinocytes, fibroblasts and melanocytes were extracted by adding 100 µL of lysis buffer RIPA (Sigma Aldrich, Missouri, USA) supplemented with a phosphatase and protease inhibitor cocktail. A 30-minute incubation of the samples on ice followed this. The sample mixture was transferred to 1.5 mL Eppendorf tubes and centrifuged for 15 minutes at 16000rpm at 4°C. Total protein dosage was further carried out using a BCA protein quantification kit (Life Technologies, California, USA). Western blotting protocol was performed as previously described. Equal protein amounts were resolved by SDS-PAGE (Life Technologies, California, USA) and transferred to a nitrocellulose membrane (IBlot gel transfer, Life Technologies, California, USA). The nitrocellulose membrane was blocked with 5% lyophilized milk or bovine serum album (BSA), followed by the addition of primary XPC antibody (1/500) incubated overnight at 4°C. Afterward, incubation with mouse HRP antibody (1/5000 diluted secondary antibody) was done for 1 hour at room temperature and following the addition of the western lightening ECL Pro ECL (Perkin Elmer), images were then directly recorded using Biorad Molecular Imager® Chemi DocTM XRS. Results were analyzed using Image Lab™ software. Glyceraldehyde-3-phosphate dehydrogenase (GAPDH) housekeeping gene was utilized to normalize the expression level of the target gene. Samples were launched in triplicates (N=3).

### Sequencing and off-target analysis

For post-genome editing analysis, wild-type and heterogeneous populations of N/TERT-2G cells were harvested, as mentioned above. DNA was extracted via QIAamp DNA mini kit (Qiagen, France) based on the manufacturer’s instructions. Using PCR primers, forward primer sequence (5’CCATTGACAGTCACCAGAGG3’) and reverse primer sequence (5’AACATAGCTGTGCCTGGACA3’), the genomic area of XPC’s exon three was amplified to yield an amplicon of 612 bases in size. Amplified amplicons were desalted and sequenced at Microsynth, France. Chromatograms were generated by the Inference of the CRISPR Edits (ICE) tool designed by Synthego to assess the genome editing efficacy based on the knockout (KO) score in a heterogeneous population versus wild-type N/TERT-2G keratinocytes. Wild-type and XPC KO N/TERT-2G, S1F/TERT-1 and Mel-ST cell lines were prepared similarly as mentioned above for final KO clones sequencing. To predict the off targets of the predesigned sgRNA utilized, CRISPOR software was utilized.

### UVB dose response

The photosensitivity of XPC KO cells was assessed and compared to wild-type cells based on the increased doses of ultraviolet B (UVB) treatment. Both wild type and XPC KO cells from each cell line and type were seeded in 6 well plates and kept until they reached 80 percent confluence. Before irradiation, they were rinsed with PBS and then exposed to escalating UVB doses. The viability of the cells was recorded 24-, 48-, and 72-hours post UVB irradiation using trypan blue assay (Thermofisher Scientific, Massachusetts, USA) based on the manufacturer’s instructions. Normalization of the data was performed by calculating at each dose the viability percentage and comparing it to control non-irradiated cells (dose 0 J/cm^2^) set as 100% viability. Samples were launched in triplicates (N=3).

### Immunofluorescence and associated microscopy

To test the repair capacity, both wild type and XPC KO cells from each cell line and type were seeded to reach 80% confluence. Afterwards, these cells were subjected to UVB irradiation. Following UVB irradiation at time 0h and after 24 hours, cells were further stained based on the protocol, which comprises the fixation of the cells using 4% paraformaldehyde and 0.2% of Triton X-100 to permeabilize the cells. 2M HCL was then utilized to fully denature the DNA double helix, enhancing the access of the antibody targeting DNA damage caused by UVB irradiation. After the saturation process, cells were incubated overnight with 1/200 primary 6-4PP antibody (Cosmo Bio, California, USA). Secondary mouse antibody 1/500 FITC (Invitrogen, California, USA) was then added the next day after several PBS washes to remove the unbound 6-4PP primary antibody. Another PBS wash removed the unbound secondary antibody to finally counterstain the DNA with Hoechst (Sigma Aldrich, Missouri, USA). With a 10X magnification, Image acquisition was done and then quantified using Cell-insight NXT. Samples were launched in triplicates (N=3). For XPC or vimentin staining, the same steps were utilized as mentioned above, excluding the step of 2M HCL incubation and by using 1/200 primary antibody targeting XPC (mouse, Santa Cruz, sc-74410 or vimentin (rabbit, Abcam, ab92547).

### Cell proliferation assay

To decipher the impact of *XPC* gene KO on the proliferation status of N/TERT-2G, S1F/TERT-1, and Mel-ST cell lines, an EDU assay was carried out. The utilization of the nucleoside analog EDU within cells has the potential to assess cellular health profile and genotoxicity. A copper-catalyzed covalent reaction between an azide and an alkyne, known as click chemistry, can be employed to quantify the integration of the nucleoside analog EDU and provide further insight into its incorporation. To do that, wild-type and XPC KO of each cell type were seeded in 6 well plates to reach 50% of confluence. Afterwards, EDU was diluted and added to the cell media for 5 hours. Subsequently, the cells were detached using trypsin, collected, and subjected to staining as per the guidelines provided by the manufacturer (Thermo Fisher Scientific, Massachusetts, USA). The analysis was carried out utilizing flow cytometry equipment (FACScan, BD LSRII flow cytometer, BD Biosciences). The post-analysis was done using flowing software (Turku Bioimaging, Finland). Samples were launched in triplicates (N=3).

### 3D Reconstructed Skin Model

A hydrogel scaffold was prepared to encapsulate cells to form a convenient matrix for all experiments. This hydrogel comprises a solution of 2.5% fibrinogen (w/v) (cat #F8630, Sigma Aldrich, France) with aprotinin (cat #A6279, Sigma Aldrich, France) and 2.5 mM of CaCl2 (cat #C8106, Sigma Aldrich, France). Wild-type and XPC KO S1F/TERT-1 Fibroblasts were suspended in 1 ml of fibrinogen to construct a human dermal equivalent. Afterwards, for the polymerization process, thrombin (cat #T4648, Sigma Aldrich, France) was added at a concentration of 1 U/ml. Then 360 microliters of this cell-laden hydrogel solution were gently and immediately added into each of the culture chambers (cat 3460, Transwell Corning, USA), which are embedded in a 12-well plate and were kept to solidify in a cell culture incubator at 37°C with 5% CO2. After 30 minutes of incubation, 2 ml of DMEM media (ref 31966047, Gibco, Life Technologies, USA) containing 10% of fetal bovine serum (Life Technologies, USA) and 1% penicillin/streptomycin (P/S) (Life Technologies, USA) were added then deposited at the bottom part of each well, and 500 microliters were added to the top of this solidified cell hydrogel matrix. The human dermal equivalent was kept in culture for 15 days. Every two days, the media was changed, 10 ng/ml of epidermal growth factor (EGF) and 80 µg/ml of vitamin C/ascorbic acid (cat A8660-5g, Sigma Aldrich, France) were supplemented to permit the maturation of the construct. Following 15 days, at passage 7, wild type and XPC KO N/TERT-2G keratinocytes and Mel-ST melanocytes were harvested. A mix of 1:40 ratio between keratinocyte and melanocyte cells was seeded onto the top of the dermal equivalent. The skin organoids were kept immersed for 9 days in the green-adapted medium. Every two days, the media was changed. On day 24, each condition was raised on the air-liquid interface. To do so, the inserts were transferred to 12 well plates to secure them, and at the same time, 4 ml of DMEM media (ref 31966047, Gibco, Life Technologies, USA) with 80 µg/ml of vitamin C/ascorbic acid (cat A8660-5g, Sigma Aldrich, France), 8 mg/ml of bovine serum albumin (cat A2153-50G, Sigma Aldrich, France) and 1% of penicillin/streptomycin (P/S) (Life Technologies, USA) were added to each condition in deep well plates so that lower surface of the insert will become in contact with the media and the upper part was deprived of media so that keratinocytes and melanocytes will differentiate. The organoids were kept in culture for 14 days, and the media was changed every 2 days. Samples were done in triplicates (N=10).

### Cytokine Assay

The LEGENDplex™ Human Inflammation Panel 1 kit (BioLegend, San Diego, USA) was utilized to measure cytokines and chemokines, which included IL-1β, IFN-α2, IFN-γ, TNF, MCP-1 (CCL2), IL-6, IL-8 (CXCL8), IL-10, IL-12p70, IL-17A, IL-18, IL-23, and IL-33. The measurements were carried out in accordance with the manufacturer’s instructions. Subsequently, samples were acquired using a FACSCanto™ II cytometer (BD Biosciences, Franklin Lakes, USA) and analyzed utilizing the online QOGNIT LEGENDplex™ program.

### Statistical Analysis

Single cell analysis were carried out by R software. GraphPad Prism v.8 was used for statistical analysis, data normalization and quantification of normality to allow the downstream selection of the respective statistical test (parametric or non-parametric) for each particular set of experiments.

## Acknowledgment

AN is supported by a fund from the doctorate school (EDISCE) at University Grenoble Alpes. WR’s contribution was funded by ANR grant PG2HEAL (ANR-18-CE17-0017) and supported by the French National Research Agency in the framework of the “Investissements d’avenir” program (ANR-15-IDEX-02).

## Authors’ Contribution

AN performed all the experiments and wrote the manuscript. ES and WR supervised the project. FK revised the manuscript. AH aided in cell culture. HR and JR aided in secretome profiling analysis. JS aided in the keratinocytes CRISPR protocol. All authors edited, read, and approved the manuscript.

